# Predicting Valley Fever Outbreaks: Novel Mechanistic Models Incorporating Climate and Ecological Interactions

**DOI:** 10.64898/2026.04.21.719799

**Authors:** Trevor Reckell, Beckett Sterner, David M. Engelthaler, Petar Jevtić

## Abstract

Coccidioidomycosis (Valley fever) is an environmentally acquired fungal infection endemic to the arid Americas, presenting a growing public health challenge as changing environmental patterns threaten to amplify exposure risks across both established and newly recognized endemic zones. Historically, forecasting efforts have relied on statistical correlations with meteorological variables. These phenomenological models often fail to capture the complex, non-linear interactions between the saprobic (environmental) and parasitic (host) life cycles of Coccidioides, particularly under non-stationary climate conditions. Here, we present a hierarchy of mechanistic Ordinary Differential Equation (ODE) models that explicitly map environmental drivers to the distinct biological stages of the fungal life cycle. We developed successive model iterations, incrementally incorporating soil moisture retention, temperature-dependent growth rates, and wildlife reservoir dynamics, and calibrated them against human case data from various regions of Arizona. We derive a time-variant environmental reproduction number and test how transmission potential fluctuates dynamically with environmental forcing. The comparative forecasting analysis, utilizing various statistical tests, information criteria, Relative Root Mean Square Error, the Diebold-Mariano test, and the Modified Diebold-Mariano, shows how the models progress. Mechanistic models based solely on continuous fungal growth perform worse than statistical baselines. By integrating climate data, we increase predictive power to a level comparable to that of the statistical model. Explicitly incorporating a wildlife reservoir as a biological amplifier significantly improves model forecasting over statistical baselines. This framework offers public health officials a biologically grounded tool to predict disease burden and guide targeted interventions responding to changing climate patterns.

## 1 Introduction

Coccidioidomycosis (Valley fever) is an environmentally driven fungal disease transmitted via inhalation of airborne *Coccidioides* spores, whose incidence is tightly coupled to climatic, ecological, and soil processes. The spores come from the fungi *Coccidioides immitis* or *Coccidioides posadasii* [1]. In the United States alone, symptomatic infections are estimated at approximately 273,000 cases annually, resulting in 23,000 hospitalizations and 900 deaths, with a direct economic burden exceeding $888 million [2, 3]. These figures substantially underestimate true incidence, as approximately 60% of infections are asymptomatic, which limits surveillance accuracy and obscures transmission dynamics [4]. Mechanistic modeling of coccidioidomycosis is challenging due to the coupling of multiple processes, including environmentally driven fungal growth in soil, host-associated parasitic dynamics, anthropogenic disturbance, and wildlife ecological interactions. The fungus *Coccidioides* that causes coccidioidomycosis has a parasitic cycle that occurs within a host and a saprobic cycle that occurs within the environment [5]. Figure 1 illustrates key structural components of *Coccidioides*, including hyphae that make up the mycelium structure and arthroconidia, which fragment from the ends of the hyphae, as well as representative ecological interactions with rodent hosts. The ecological context includes rodent hosts such as kangaroo rats (*Dipodomys merriami*) and woodrats (*Neotoma lepida*), which are known reservoirs. Both species are prevalent in Arizona and commonly infected with coccidioidomycosis[6][7]. Despite its prevalence in arid regions of the Americas, particularly in the Southwestern United States, mechanistic modeling of Valley fever has been limited. To address this gap, we develop a hierarchy of mechanistic models that explicitly couple soil fungal dynamics, environmental forcing, and host exposure pathways. These steps enable systematic analysis of transmission drivers and emergent disease dynamics.

**Figure 1:**
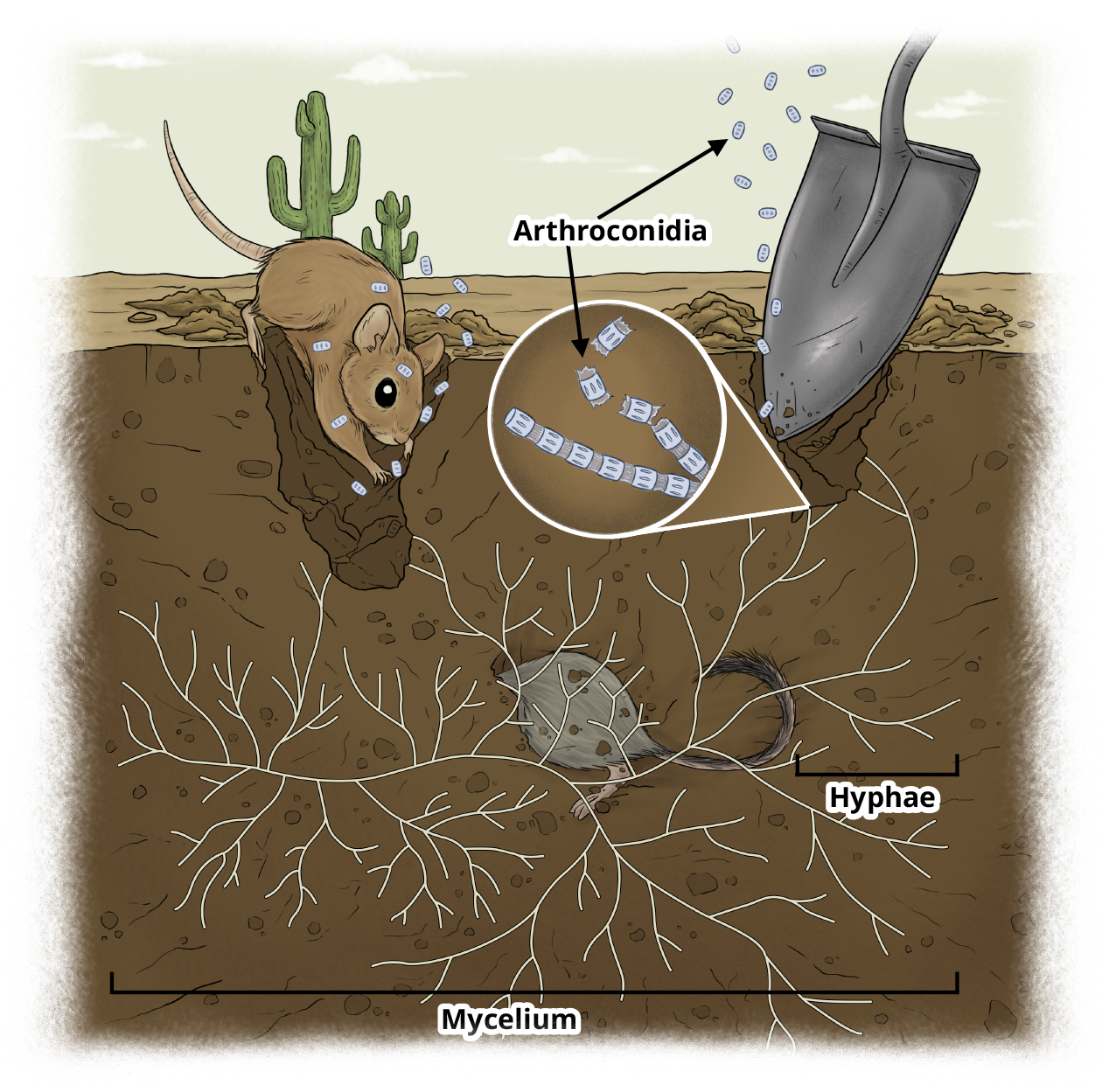
A diagram of *Coccidioides* in the environment. Arthroconidia are asexual spores produced by certain fungi. Note, this diagram is not to scale as the mycelium, hyphae, and arthroconidia of *Coccidioides* are all between 2-5 micrometers [8].

### 1.1 Past modeling efforts

Mechanistic modeling efforts for coccidioidomycosis remain limited, with most prior work relying on statistical or correlational analyses linking incidence to climatic variables citecomrie2005climate [9]. One exception is the agent-based model developed by Anderson [10], which incorporates environmental and demographic heterogeneity. However, broader efforts to create deterministic models that capture the underlying biological and environmental processes have been lacking. These limitations underscore the need for deterministic, process-based models capable of systematically exploring the biological and environmental mechanisms underlying the observed outbreak patterns.

### 1.2 Coccidioides Substrate

The dimorphic fungal pathogens *C. immitis* and *C. posadasii* utilize a specialized saprophytic nutritional strategy during their saprophytic life cycle. These fungi are adapted to decompose animal-derived organic matter, particularly small mammal remains [11, 7]. Their genomes are enriched with genes for enzymes that break down animal-derived proteins and fats while being notably deficient in genes required for degrading plant material [7, 12]. Consequently, these *Coccidioides* species do not effectively break down dead plants. While the fungus only survives on specific soil chemistries, notably arid, alkaline soils with high salinity and elevated levels of minerals like boron, calcium, sodium, and magnesium, these abiotic factors do not serve as a primary energy source [13]. It is hypothesized that this harsh geochemical environment functions to create a selective niche by inhibiting microbial competitors, allowing *Coccidioides* to thrive [14]. Laboratory experiments confirm that the fungus can persist and grow using soil as a sole source of nutrients, but this likely represents a survival state [13]. The leading hypothesis is that robust, reproductive growth that produces the infectious arthroconidia is fueled by the nutrient windfall provided by an animal carcass [11]. In summary, *Coccidioides* breaks down the proteins of carcasses of dead mammals that had previously been infected, persists on ambient soil nutrients, and inhabits a specific mineralogical environment that facilitates its unique life cycle. This substrate specificity motivates the inclusion of organic matter and wildlife derived organic matter as a key driver of environmental fungal growth in our modeling framework.

## 1.3 Fungal Models

A foundational component of any mechanistic Valley fever model is the life cycle of the *Coccidioides* fungus itself. The growth of filamentous fungi has been described using partial differential equation (PDE) models that account for the density of different fungal structures like hyphae [15]. While these models provide a high level of detail, they can lack validation with empirical fungal data and may be too complex for direct integration into a larger epidemiological system [15]. Methods to simplify these PDE systems into more manageable ordinary differential equation (ODE) frameworks have been proposed [16]. Unlike in most PDE models, in our ODE models proposed we will operate under the assumption of spatial homogeneity for all compartments.

Simpler models that capture the essential dynamics are more practical. Logarithmic models linking fungal growth directly to environmental factors like temperature and water activity have been developed, offering a more straightforward approach [17]. Although we initially looked at the paper by Sautour et al. [17], we had difficulties recreating their results and preferred a smoother function in extreme conditions. Thus, we use the Sautour et al. paper as inspiration for a starting point for our functions later that describe the effect temperature and soil moisture has on fungal growth in the saprobic cycle. These effects play a role in the *Coccidioides* life cycle: the growth of the mycelium (an interconnected network of hyphae), the production of arthroconidia (spores), and the establishment of new fungal colonies [5].

### 1.4 Environmental Drivers: Temperature and Soil Moisture

The transmission of Valley fever is intrinsically linked to environmental conditions governing its saprobic cycle. The fungus is uniquely adapted to extreme weather fluctuations: heavy precipitation increases soil moisture and facilitates nutrient acquisition and rapid hyphae growth in the soil, while subsequent severe dry periods act as a biological trigger, forcing the fungus to segment into durable, infectious arthroconidia [18]. It is this specific mechanism of desiccation-induced aerosolization, rather than simple continuous growth, that drives human exposure. Therefore, precise mechanistic modeling of the sequential dynamics of soil moisture and temperature is critical. Once formed, these spores become airborne through wind or other soil disturbances. However, before being dispersed in the air, the arthroconidia spores must be brought up from under the soil surface through physical mechanisms, such as human or wildlife soil disturbances. Once at the surface, weather dependence takes over, and weather patterns have been a key focus in modeling other climate-sensitive diseases, such as malaria [19],[20]. Extensive work has been done to create temperature- and rainfall-dependent models for vector-borne diseases, which can serve as an inspiration for our framework. These models often use functions to describe how temperature and humidity affect vector and parasite life cycles. Though Coccidioidomycosis is not a vector-borne disease, wildlife still spreads and acts as a reservoir for the fungus, so we can learn from many of the climate models utilized in vector-borne diseases. Following this approach, we will incorporate functions that describe the multiplicative effects of temperature and soil moisture on the growth rates of both hyphae and arthroconidia, using optimal temperature (*T*_*opt*_) and soil moisture (*S*_*opt*_) parameters to define these environmental windows for fungal proliferation.

### 1.5 The Role of Wildlife

While wind has been hypothesized as the primary factor in the dispersal of *Coccidioides* spores, the activity of burrowing animals is another crucial mechanism for bringing spores to the surface [21]. The arid environments of the United States and Mexico are home to numerous burrowing animals, including various species of squirrels, pocket mice, and woodrats. The burrowing activity of these semi-fossorial vertebrates is a critical ecological process that can significantly influence spore aerosolization [22]. The genetic diversity and distinct populations of *Coccidioides* across different geographic locations suggest that while dispersal occurs, possibly from wildlife movement, the fungus also maintains localized environmental niches that are frequently disturbed [23].

To account for this, our more advanced models will incorporate a reservoir compartment representing the population dynamics of these key burrowing animals. We will refer to organisms in this compartment as “wildlife,” representing the population size of relevant wildlife, such as burrowing animals. We also use the term “reservoir” for the compartment, as the wildlife contributes to the maintenance of *Coccidioides* in the wild. The life cycle and social structures of relevant burrowing species, such as the bushy-tailed woodrat (*Neotoma cinerea*), have been well-documented and can inform the structure of models [24]. However, more in-depth reservoir dynamics will be a part of future work. We will model the wildlife population’s breeding season as a function of temperature, reflecting the spring and fall breeding patterns common to most mammals in the region. The death of these wildlife, in turn, contributes to the organic matter in the soil that *Coccidioides* can feed on, creating a feedback loop that provides food for the fungus and completes the ecological cycle.

To go from statistical forecasting to utilizing the underlying biological mechanisms, we developed a hierarchy of ODE models that explicitly couple the environmental dynamics of *Coccidioides* with human epidemiology. By systematically introducing soil moisture, temperature-dependent fungal growth rates, and wildlife reservoir dynamics, we tested the relative impact of these ecological drivers on disease transmission. Our comparative analysis methodology against historical data reveals that explicitly modeling the saprobic cycle, particularly the extreme weather fluctuations that trigger arthroconidia production (which can late be aerosolized), is critical for capturing the non-linear dynamics of Valley fever outbreaks. We demonstrate that incorporating the substantial asymptomatic fraction of the population alters the inferred transmission parameters, providing a more realistic estimate of the burden of the disease. The mechanistic framework we propose offers a scalable and biologically grounded alternative to traditional correlational models, providing public health officials with an improved tool to anticipate high-risk periods in an era of shifting climates.

From a modeling perspective, our approach differs from prior work in multiple crucial ways. We explicitly represent environmental fungal dynamics as a coupled subsystem, incorporate climate-dependent forcing functions, and construct a hierarchical modeling framework that enables mathematical and analytical investigation of key epidemiological quantities, including the basic reproduction number. In addition, the more advanced models involve wildlife as both a direct driver of environmental fungal growth and a potential reservoir host, further linking ecological and epidemiological dynamics.

## 2 Modeling Framework

To introduce the modeling framework, we present a series of increasingly complex mechanistic ODE models that represent the potential effects of different factors on the etiology of Valley fever from soil growth to human infection (Figure 2). Since *Coccidioides* has a parasitic cycle that occurs within an animal host and a saprobic cycle that occurs within the environment, we break each model into two distinct parts: a Fungal Model and a Human Exposure Model. The Fungal Models focus on the saprobic cycle and spread in the environment, while the Human Exposure Models focus on human exposure and infection. Figure 2 illustrates the conceptual structure of the overall framework.

**Figure 2:**
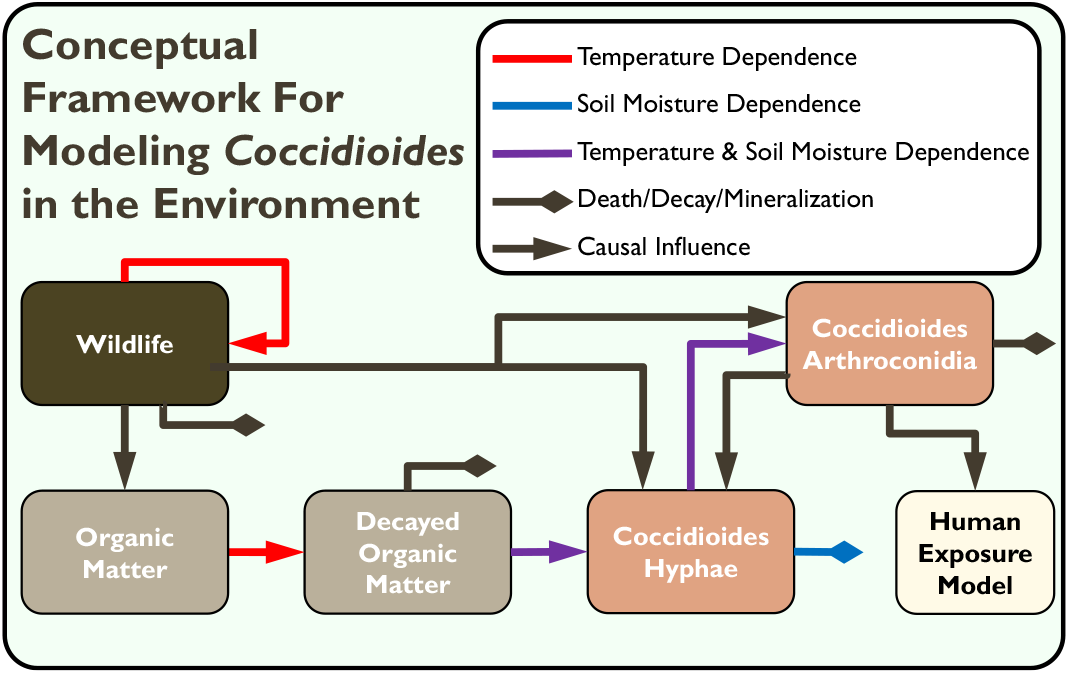
Visual illustration of full conceptual framework for modeling Valley fever dynamics corresponding to the most complex models (Model 4a/4b) introduced in this paper. Colored boxes show key model variables and arrows show causal interactions. Colored lines indicate when causal interactions include dependence on external environmental conditions. Models 1-3 are distinguished from Models 4a/4b by excluding key variables and interactions.

### 2.1 Simple Fungal Growth and Human Exposure: Model 1

As a foundation for more complex and realistic models, we first introduce a system of equations for saprobic fungal growth that represents the dependence of hyphae on decayed organic material. We also introduce a simple Susceptible-Infected-Recovered (SIR) system of equations for human disease cases, where the rate of new infections is proportional to the amount of hyphae in the system.

#### 2.1.1 Fungal Model 1

The simplest fungal model we have laid out has two variables: decayed organic matter biomass (*D*) and hyphae biomass of *Coccidioides* (*H*). The parameter *O* represents new decayed organic matter which *Coccidioides* hyphae cells consume. The parameter *μ*_*H*_ represents the rate of uptake of decayed organic matter by the hyphae. Lastly, *δ*_*D*_ represents the decay rate of the decayed organic material to a point where nutrients from the material can no longer be used. Then we describe the dynamics of the decomposed organic matter compartment as

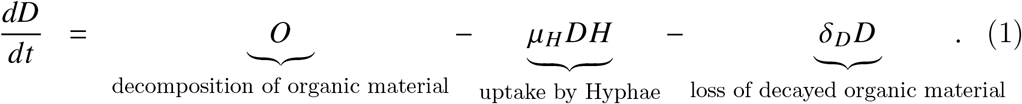

For the hyphae compartment, the parameter *γ*_*H*_ represents the growth rate of hyphae, *H*_*max*_ represents the maximum hyphae population in a region given non-decayed organic matter-related nutrients and competitive restrictions, and *δ*_*H*_ represents the death rate of hyphae. The parameter *H*_*max*_ contributes a logistic model behavior to the hyphae compartment and mainly serves to limit population growth. Logistic models were conceived in the context of human population dynamics [25], but they have also been applied to the growth of other organisms, including the growth of hyphae [26]. Then hyphae biology takes the form of

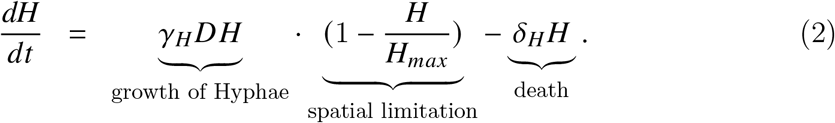

Thus, Fungal Model 1 is simply the decayed organic matter compartment in Equation 1 and the hyphae of *Coccidioides* compartment in Equation 2.

#### 2.1.2 Human Exposure Model 1

For the human exposure process, we start with a simple open population SIR model. The variables or compartments of susceptible humans (*S*), infected humans (*I*), and recovered humans (*R*) come from the original SIR model papers by Kermack and McKendrick [27, 28, 29]. A crucial difference here is that people are infected based on interaction with the fungus (in Model 1 via hyphae, but in later models arthroconidia) rather than infected humans. The combined total population (*N*) is defined as the sum of all human compartments. The parameters, *α*_*h*_ and *ω*, describe the birth/immigration and death/emigration of humans, respectively. There is also the parameter *c* representing the density-dependent death rate. Behind all the human exposure models in this paper is a human population model that is a variant of the logistic model [30]. The human population model chosen is rearranged algebraically to better fit multiple compartments for humans [31] using the parameter 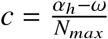, where *N*_*max*_ is the carrying capacity of humans in a given region. Thus the overall human population logistic equation:

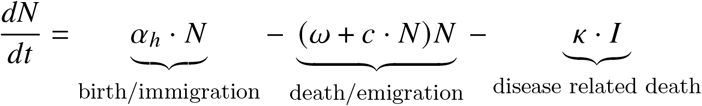

The parameter *ε* controls the rate of infection of Valley fever relative to the hyphae. In later models, the *ε* parameter describes the rate of infection relative to arthroconidia. The parameter *ϱ* is the rate of recovery, and the parameter *κ* is the Valley fever-specific death rate. Note the absence of re-susceptibility, since this occurs so rarely and people normally have lifelong immunity [32, 33]. The Human Exposure Model 1 is then laid out as

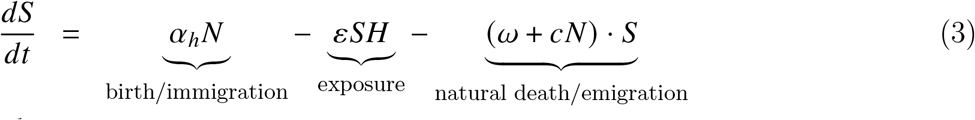

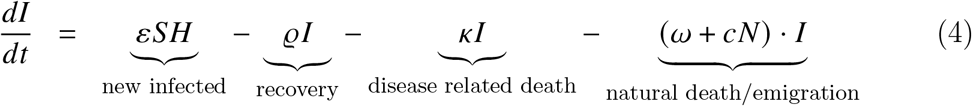

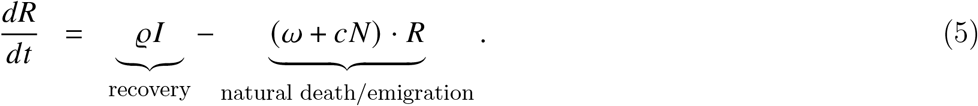

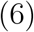

### 2.2 Arthroconidia Production and Exposure: Model 2

An obvious gap in the fungal component of Model 1 is the importance of arthroconidia in the infection of new hosts. Rather than implicitly assuming the number of newly infected individuals are proportional to the amount of hyphae, we can explicitly represent the production of arthroconidia from hyphae and exposure to arthroconidia as the key mechanism for infection. Additionally, arthroconidia play an important role in hyphae growth when infected hosts die and found new colonies.

#### 2.2.1 Fungal Model 2: Relations of hyphae and arthroconidia

Arthroconidia biomass (*A*) is a new variable with a growth rate of *γ*_*A*_ and a death rate of *δ*_*A*_. The rate of arthroconidia spreading and successfully forming a new colony, *φ*_*A*_, is both a mechanism of arthroconidia loss and hyphae growth through the term *φ*_*A*_ *A*. The hyphae compartment is similar to Fungus Model 1, now with additional growth coming from arthroconidia colonizing new areas.

The primary food for *Coccidioides* is organic matter, specifically dead animals, which we now classify as a new variable in its own compartment as (*O*). The amount of new organic matter coming in is defined as the parameter Π. The decay rate of the organic matter is defined as *δ*_*O*_. The decomposed organic matter is still defined as (*D*) and remains similar to Fungus Model 1, with the small change that the decomposition of organic matter now takes the form of *δ*_*O*_*O*.

Together, with the biological interactions of arthroconidia, the compartments of Fungal Model 2 are laid out as:

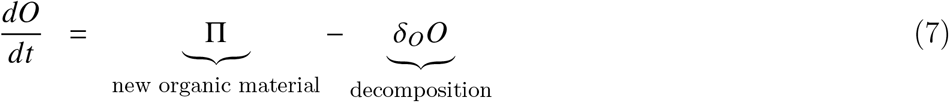

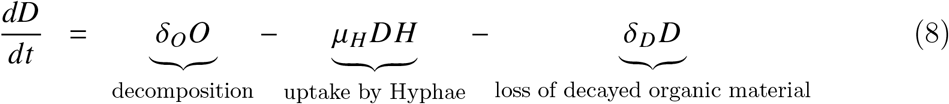

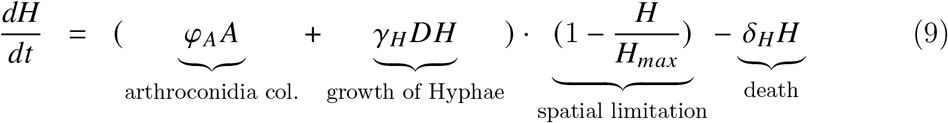

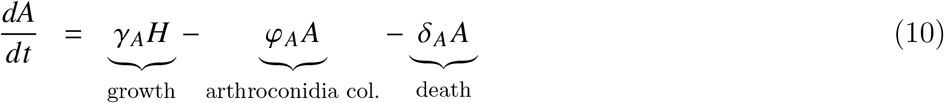

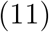

Some assumptions being implicitly made in the model layout are that all organic and decomposed matter decay and stay in the system and that only *Coccidioides* remove decomposed matter. Additionally, we are assuming that this system is at an appropriate temperature and water level for the asexual reproduction of arthroconidia [34] depending on the set level of *γ*_*A*_. However, *Coccidioides* is mainly observed to reproduce asexually in all environments [35]. Fungi typically produce spores at a much higher rate during asexual reproduction compared to sexual reproduction.

#### 2.2.2 Human Exposure Model 2

Human Exposure Model 2 contains the additional variable of the exposed human compartment (*E*), and the parameter *ψ*, which is the rate of becoming symptomatic or diagnosable. The addition of the exposed compartment for the latent period of a disease was popularized in a book by Bailey [36]. Humans are not susceptible to Valley fever from other infected individuals, except in cases of organ transplants or congenital transmission [37]. Therefore, infected humans are those with diagnosable symptoms, and exposed humans are those with the fungal infection but are undiagnosed and show no clear symptom combination of coccidioidomycosis. With these changes, Human Exposure Model 2 becomes

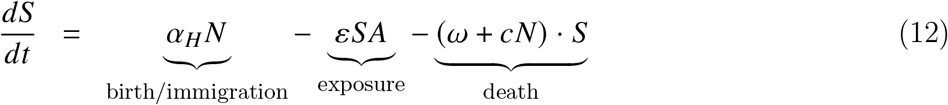

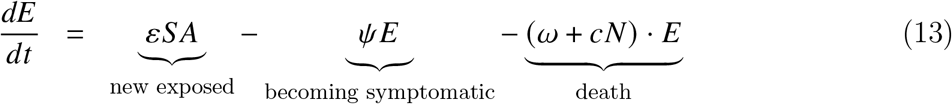

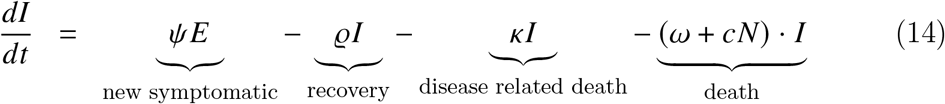

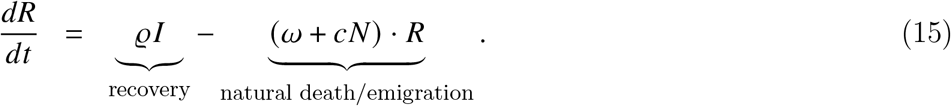

### 2.3 Environmental Factors: Model 3

Environmental factors are crucial for the growth of fungus hyphae and arthroconidia. The two most important factors for these are temperature and soil moisture [18].

Model 3 incorporates these effects into the fungal dynamics but uses the same human exposure system as Model 2 (2.2.2).

#### 2.3.1 Temperature and Soil Moisture Effect on Fungal Growth: Fungal Model 3

Fungal Model 3 uses two additional data sets, temperature, *T*, and soil moisture, *S*_*m*_. The compartments of Fungal Model 3 remain the same as Fungal Model 2, but the effects of temperature and soil moisture are new. We assume that temperatures stay within a range which organic decay occurs, and the parameter *T*_*decay*_ is the inflection point above which organic matter decomposes faster and below which it decomposes slower. Thus the simple Hill function for temperature dependent organic decay is 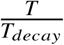.

Fungal growth rates are multiplied by functions of environmental variables that describe their effects on fungal growth rates. Temperature and soil moisture are entered as time-dependent forcing terms derived from empirical data and incorporated via a piecewise cubic Hermite interpolating polynomial (PCHIP) of the observed environmental time series. We define the function for the multiplicative effect temperature has on hyphae as 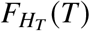 and the function for the effect temperature has on arthroconidia as 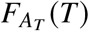. Both functions utilize the following parameters: *α*_*T*_, a scalar for the effect of temperature above optimal temperature; *β*_*T*_, a scalar for the effect of temperature below optimal temperature; and *T*_*opt*_, the optimal temperature. The values and ranges for *α*_*T*_, *β*_*T*_, and *T*_*opt*_ are independent in the respective functions 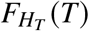 and 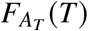. Based on previous work [38, 39], the overall structure of 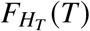 and 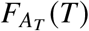 then takes the form of a piecewise Gaussian function,

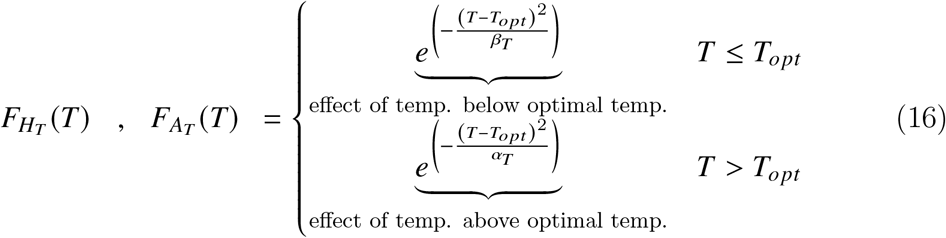

Similar to 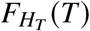 and 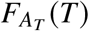, the multiplicative effect that soil moisture has on hyphae 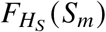 and arthroconidia 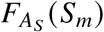, utilize the parameters *S*_*opt*_, the optimal soil moisture level for growth, *β*_*S*_, scalar for effect of soil moisture below optimal soil moisture level, and *α*_*T*_, scalar for effect of soil moisture above optimal soil moisture level. Again, it should be noted that the ranges and values are independent for parameters (*S*_*opt*_, *β*_*S*_, *α*_*S*_) in 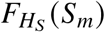 and 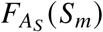. The effects simply take the same structural form, which once again is a piecewise Gaussian function. Other work has used the same type of functions for the effects of “water activity” as many papers call it [40, 41, 42], which we call soil moisture. Then 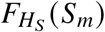 and 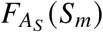 are

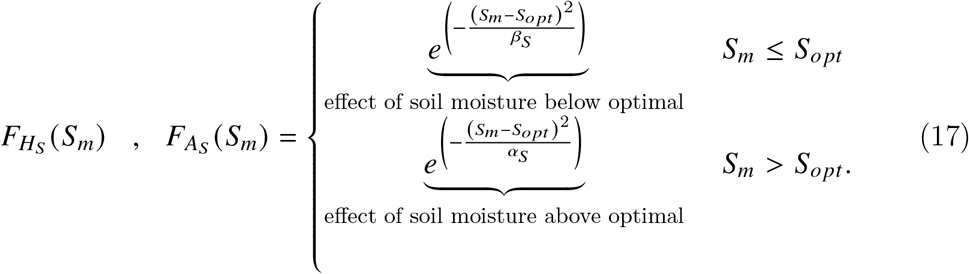

Note *T*_*opt*_, *S*_*opt*_, *β*_*T*_, *β*_*S*_, *α*_*T*_ and *α*_*S*_ are all greater than 0. In Equation 16 and Equation 17, *T*_*opt*_ and *S*_*opt*_ would be constants if lab data providing optimal temperature and soil moisture levels for Coccidioides was available. Instead, *T*_*opt*_ and *S*_*opt*_ are fit parameters due to the lack of such data.

The combined effect of soil moisture and temperature on hyphae and arthroconidia is simply the multiplicative effect of each respectively. Both older [40] and newer [42] works show how the multiplicative effect of temperature and soil moisture effect on fungus is optimal. Thus *F*_*H*_ (*T, S*_*m*_) in Equation 18 and *F*_*A*_ (*T, S*_*m*_) Equation 19 are defined as

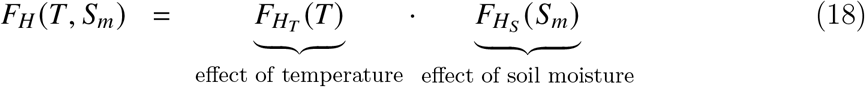

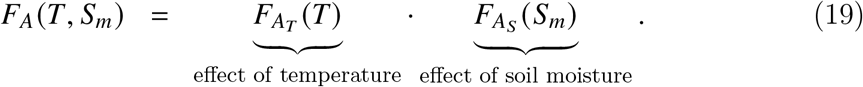

The overall dynamics of Equations 18 and 19 are similar to that shown in Figure 3, which displays an example of 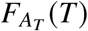. Parameters *β*_*T*_, *α*_*T*_, *β*_*S*_, *α*_*S*_ are fit parameters. However, as mentioned before, *T*_*opt*_,*S*_*opt*_,*β*_*T*_, *α*_*T*_, *β*_*S*_ and *α*_*S*_ are different if they are for hyphae (as represented in Equation 18) or for arthroconidia (as represented in Equation 19). The best practice would be to have all these parameters fitted to the growth of the specific *Coccidioides* fungus (hyphae/arthroconidia) being modeled. In the absence of this data, the parameters can be used in fitting to the Infected human data. With this Fungal Model 3, the attached Human Exposure model 2 forms the overall Model 3.

**Figure 3:**
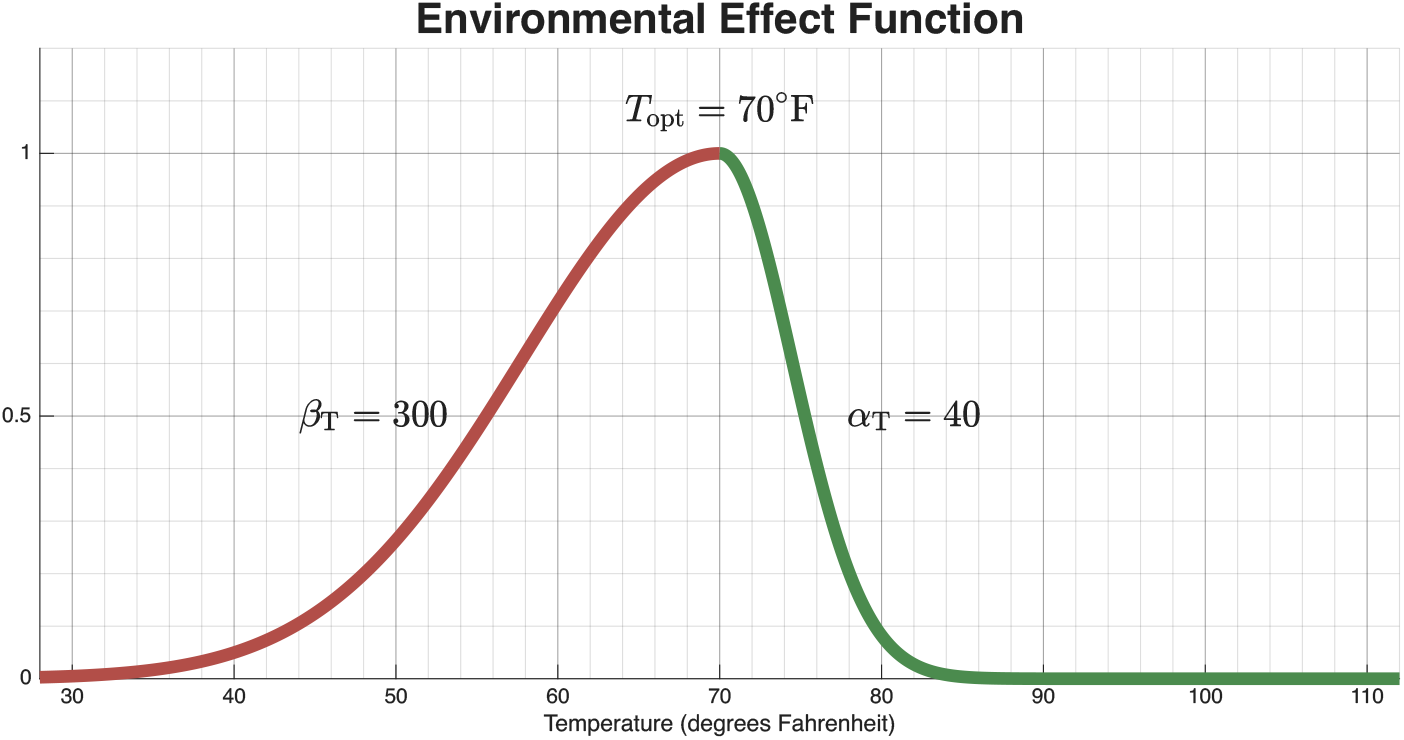
A figure showing the dynamics of 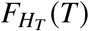, where temperature in Fahrenheit is the x-axis. The dynamics of 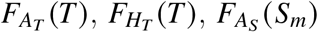, and 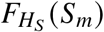 all follow this same dynamic and all have a range of [0,1] for the stated parameter ranges. Since the combined effects of soil moisture and temperature are multiplicative, the values range from 0 to 1 for Equations 18 and 19, and also follow this same dynamic.

Then the Equations 18 and 19 can be used in the hyphae and arthroconidia compartments, respectively, giving us Fungal Model 3 in the form of

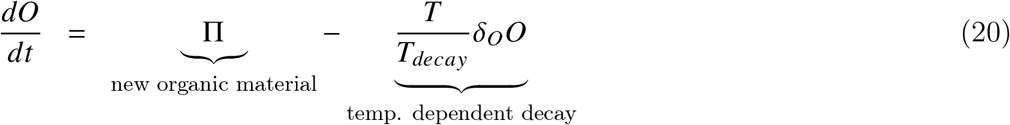

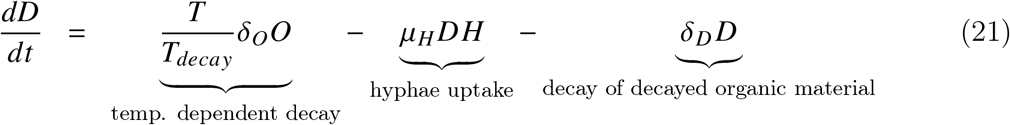

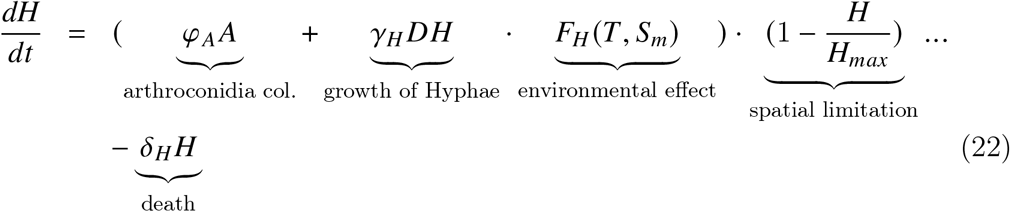

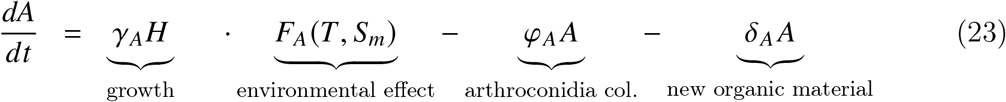

### 2.4 Wildlife Reservoir: Model 4a

Numerous animals get infected with and spread *Coccidioides* fungal spores in Arizona, including various coyotes, wolves, mountain lions, bobcats, javelinas, mule deer, deer, elk, squirrels, mice, chipmunks, rats, rabbits, badgers, shrews, skunks, bears, bats, and even some reptiles like gila monsters [43, 44, 45]. Many animals that contract Valley fever show no symptoms or only mild signs of illness [46, 45]. For simulation purposes, we will group all animals that carry *Coccidioides* into a single wildlife compartment (*W*). As stated before, we use the term “reservoir” to reflect the wildlife compartment’s role in maintaining *Coccidioides* in the environment [43]. In future work, the wildlife compartment could be expanded to represent the dynamics of a particular wildlife species or group, adding a third submodel to the overall framework.

#### 2.4.1 Fungal Model 4a

When laying out the wildlife compartment, we need to take into account the seasonality of population spikes. We adopt a phenomenological approach for wildlife population growth. Most small mammals, which form a significant part of the reservoir population, have population spikes relative to key climatic variables, since reproduction is an energetically costly activity timed to coincide with favorable environmental windows [47]. Both precipitation (which dictates food availability) and ambient temperature are established drivers of seasonal breeding in mammals. We allow for population growth to spike within certain temperature ranges, a relationship observed in some arid-zone species where temperature acts as a primary trigger for breeding [47, 48]. Then the wildlife population growth function, *F*_*wpg*_ shown in Equation 24, has the following parameters: *T*_*hs*_, the heat stress temperature for population growth, and *T*_*cs*_, the cold stress temperature for population growth. Together, these parameters define the temperature range which population growth can occur for the wildlife compartment *W*. The wildlife population growth function is defined as

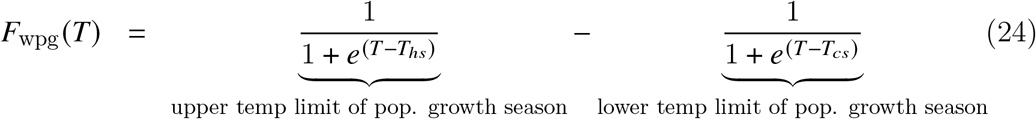

The population growth rate of wildlife is defined as parameter *β* and the death rate of wildlife is defined as parameter *δ*_*W*_. An example of how this wildlife population growth function dynamics look can be seen in Figure 4. Note that the wildlife compartment, as defined in Function 25, is a simplification of realistic wildlife dynamics. The wildlife compartment lacks a carrying capacity and realistically is dependent on many other factors not incorporated. However, with the current lack of any reliable data set or proxy to measure wildlife that spread Valley fever in the regions we are looking, a simplified compartment with minimal parameters that still changes seasonally was the goal. If wildlife data pertinent to *Coccidioides* becomes available, a more mechanistic population growth model can be implemented.

**Figure 4:**
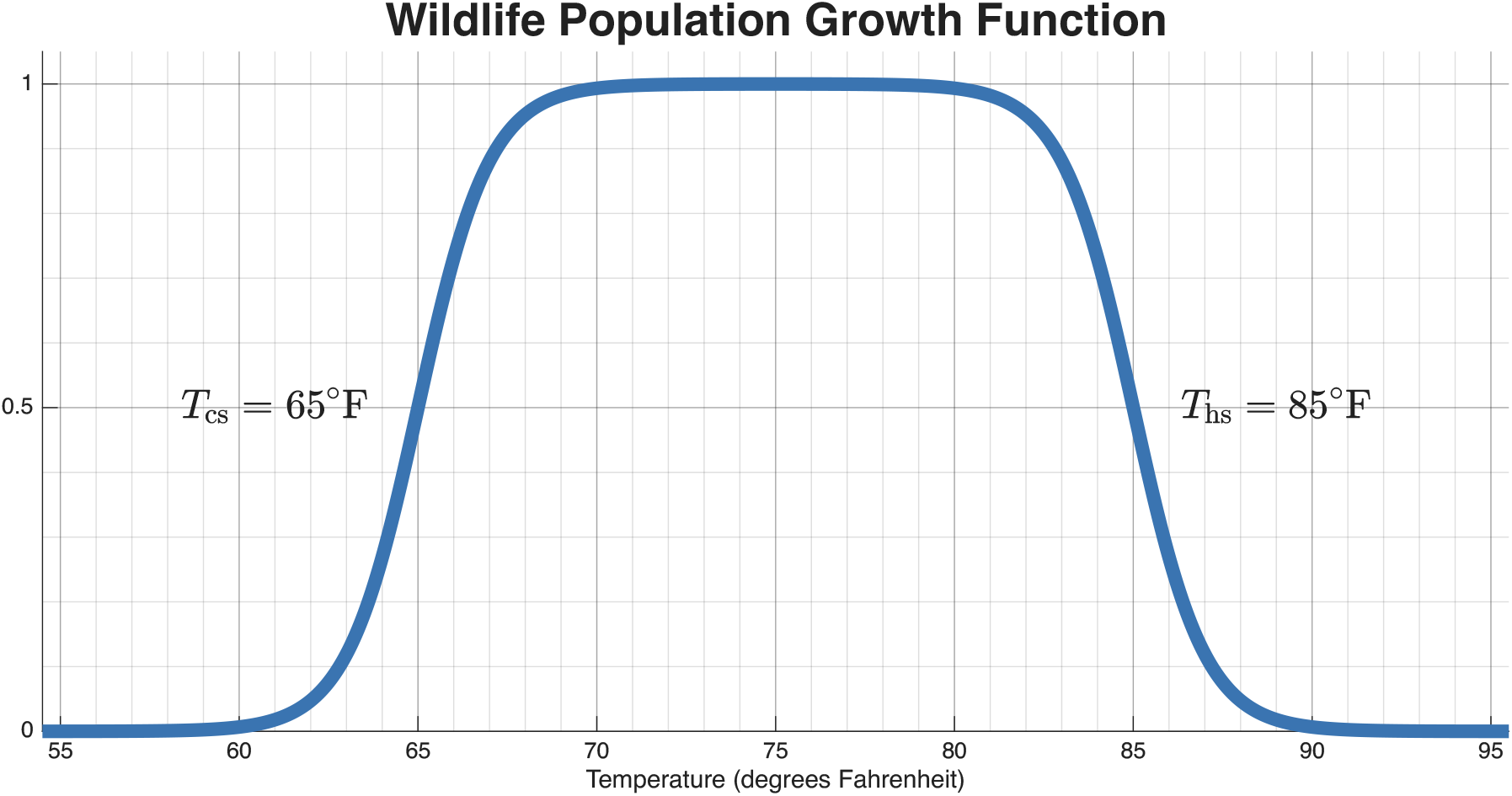
This figure is an example of the wildlife population growth function defined in Equation 24, with the x-axis of temperature in Fahrenheit. The function has a range of [0,1] for the stated parameter range.

With wildlife incorporated into the model, a new dynamic comes in the form of the *α* parameter, which is the rate of wildlife spread of arthroconidia. The parameter *α*, multiplied by the wildlife population (*W*) and the arthroconidia *A*, can be seen in the hyphae compartment as wildlife contribute to creating new colonies of Coccidioides. Wildlife defecation is added to the decayed organic matter compartment under the term *σW*, where *σ* is the defecation rate. Defecation is biologically important, as the localized accumulation of animal droppings in rodent burrows provides the highly concentrated microenvironments required for *Coccidioides* to proliferate amidst broader desert nutrient scarcity [11, 49]. Another new dynamic is from droughts, coming in the form of drought effect *F*_*dr*_. The parameters within *F*_*dr*_ are *S*_*ds*_, the drought stress soil moisture level, *T*_*ds*_, the drought stress temperature level, and *χ*, the drought effect scalar. Thus, Fungal Model 4a is

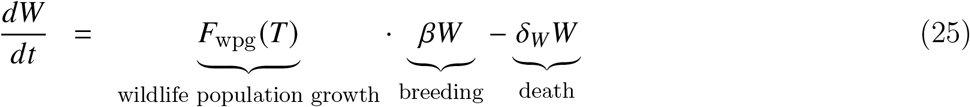

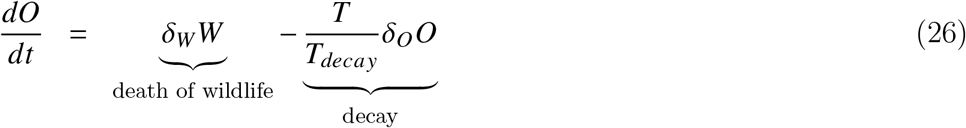

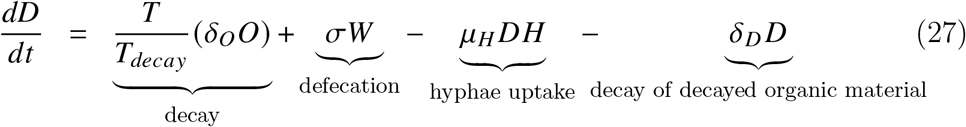

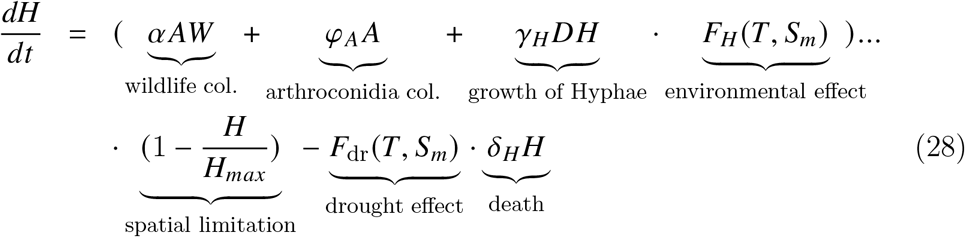

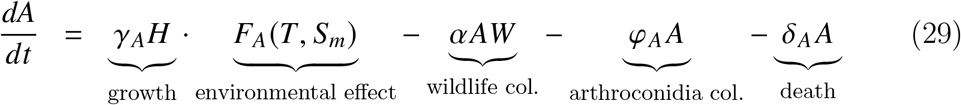

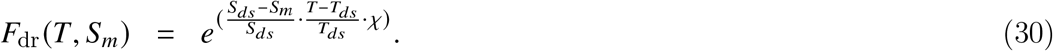

### 2.5 Asymptomatic Humans and More Mechanistic Organic Decay: Model 4b

Several mechanisms, including specifically simplifying organic decay and excluding an asymptomatic human infected compartment, have been carried through Models 2, 3, and 4. There are relatively easy additions to make to the Fungal Models and Human Exposure models to make the organic decay more mechanistic and include an asymptomatic human infected compartment, thus making an overall more mechanistic model.

#### 2.5.1 Fungal Model 4b

A simple yet more mechanistic model for the decay of organic material than what is described in previous Fungal Models is called the *Q*_10_ model, in reference to the rate of organic decay for every ten degrees Celsius change. The *Q*_10_ model was first described in 1884 [50]. The *Q*_10_ model has faults [51], but it has very few parameters compared to other organic decay functions and is considerably more mechanistic than our previous efforts, so it is a meaningful improvement. The parameter *Q*_10_ is the temperature coefficient for organic decay. Then *R*_1_ is the rate of the process at temperature *T*_1_ and *R*_2_ is the rate of the process at temperature *T*_2_. The terms *T*_1_ and *T*_2_ are the two different temperatures, usually in Celsius but can easily be adapted to Fahrenheit. Then *Q*_10_ is described as

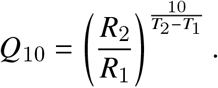

For our purposes, we don’t have *R*_1_ and *R*_2_ values for relevant organic matter and temperatures. However, we do have a clear range. The parameter *k* is the decay rate at temperature *T*, where *T* is the temperature for which we predict the decay rate. The parameter *k*_*ref*_ is the rate of decay measured at a reference temperature. Then *Q*_10_ is now temperature coefficient for the organic decay process. The parameter *T*_*ref*_ is the same reference temperature used to find *k*_*re f*_. Thus, giving us

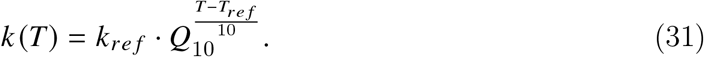

Then Fungal Model 4b can be written as

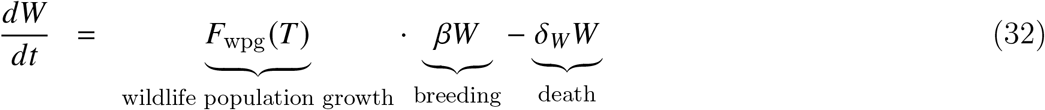

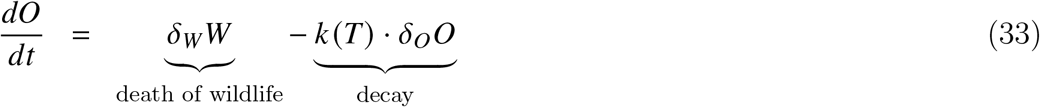

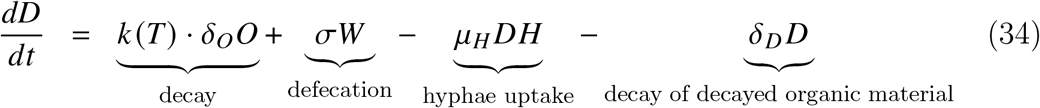

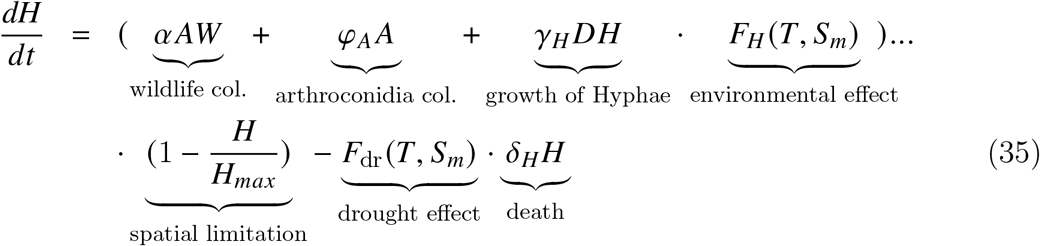

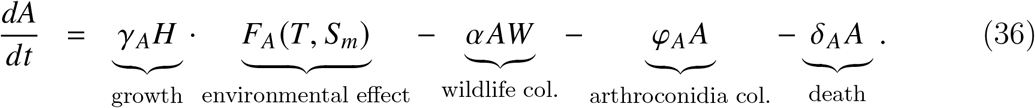

#### 2.5.2 Human Exposure Model 3

The Human Exposure Model 3 incorporates a new compartment for Asymptomatic Humans, designated (*A*_*H*_). Approximately 60% of coccidioidomycosis infections are estimated to be asymptomatic [4]. The Asymptomatic Humans definition we will adhere to is all unreported/unidentified infections, including both truly asymptomatic individuals and symptomatic patients who are misdiagnosed or never interact with the healthcare system. The extra compartment *A*_*H*_ will allow the model to capture this large amount of asymptomatic humans. The way we have formulated the *A*_*H*_ compartment means it does not offer the benefit of producing different dynamics for the infected compartment. The rate of new asymptomatic utilizes a rate of *ψ*_*A*_ as opposed to the now rate of becoming symptomatic of *ψ*_*I*_. Asymptomatic humans, the same as symptomatic, can spread Valley fever back into the soil via human burial, organ transplantation, or congenital transmission. However, these types of transmission methods are considered to be very rare, and humans are considered “dead-end hosts” for this disease. Together Fungal Model 4b and Human Exposure Model 3 form the overall Model 4b. The complete equations of Human Exposure Model 3 are,

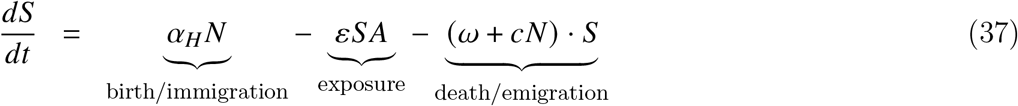

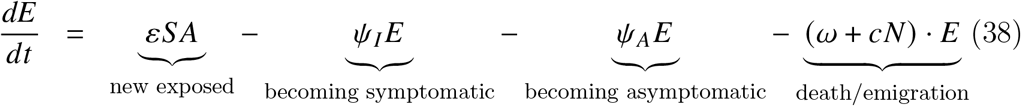

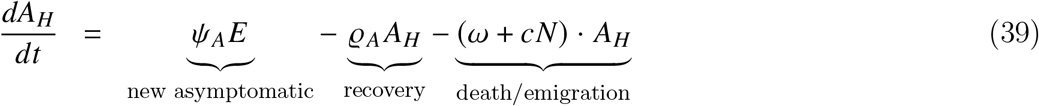

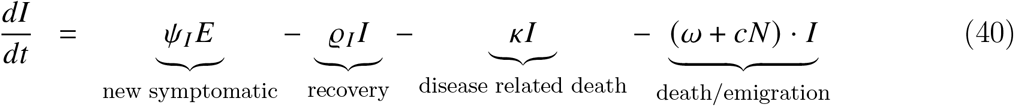

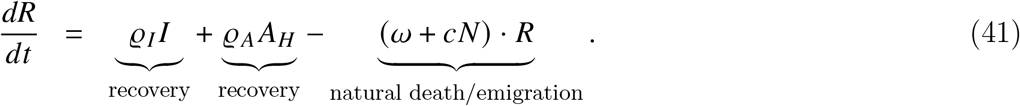

To ensure the biological validity and mathematical well-posedness of our framework for all the ODE model propose, we establish that all state variables representing physical populations (fungal biomass, organic matter, wildlife, and humans) remain non-negative for all time *t* ≥ 0 given non-negative initial conditions. Furthermore, we verify that in our stated modeling conditions, these populations do not exhibit unbounded growth. The complete mathematical proofs demonstrating positivity and boundedness for the state variables are detailed in the Supplementary Information.

### 2.6 Parameter Overview

We recognize that there are a large number of parameters introduced throughout the ODE models we have outlined, so we present Table 1 to recap the parameters used in Model 1–4b. While Table 1 provides a high-level summary of the parameters utilized across the modeling framework, the complete list of parameter constraints, bounds, and the fitted values resulting from the particle swarm optimization for each region can be found in the Supplementary Information.

**Table 1:** Parameters used in the mechanistic non-statistical models (Models 1–4b). A generic description and the specific models utilizing each parameter are provided.

| Parameter | Description | Models Used |
| --- | --- | --- |
| *Fungal Life Cycle & Environmental Drivers* |  |  |
| $O$ | Rate of new decayed organic matter input | Model 1 |
| $\Pi$ | Rate of new organic matter | Models 2–4b |
| $\mu_H$ | Rate of organic matter uptake by hyphae | Models 1–4b |
| $\delta_D$ | Decay rate of decomposed organic matter (to a state no longer usable by *Coccidioides*) | Models 1–4b |
| $\delta_O$ | Decay rate of organic matter | Models 2–4b |
| $\gamma_H$ | Growth rate of hyphae at optimal temperature and soil moisture | Models 1–4b |
| $H_{max}$ | Carrying capacity of hyphae | Models 1–4b |
| $\delta_H$ | Death rate of hyphae | Models 1–4b |
| $\gamma_A$ | Growth rate of arthroconidia at optimal temperature and soil moisture | Models 2–4b |
| $\delta_A$ | Death rate of arthroconidia | Models 2–4b |
| $\varphi_A$ | Rate of arthroconidia based colonization/spread | Models 2–4b |
| $T_{decay}$ | Temperature inflection point for organic decay | Models 3, 4a |
| $T_{opt}$ | Optimal temperature for growth (fit separately for hyphae/arthroconidia) | Models 3, 4a, 4b |
| $\alpha_T$ | Scalar for temperature effect above $T_{opt}$ (fit separately for hyphae/arthroconidia) | Models 3, 4a, 4b |
| $\beta_T$ | Scalar for temperature effect below $T_{opt}$ (fit separately for hyphae/arthroconidia) | Models 3, 4a, 4b |
| $S_{opt}$ | Optimal soil moisture for growth (fit separately for Hyphae/Arthroconidia) | Models 3, 4a, 4b |
| $\alpha_S$ | Scalar for soil moisture effect above $S_{opt}$ (fit separately for hyphae/arthroconidia) | Models 3, 4a, 4b |
| $\beta_S$ | Scalar for soil moisture effect below $S_{opt}$ (fit separately for hyphae/arthroconidia) | Models 3, 4a, 4b |

| Parameter | Description | Models |
| --- | --- | --- |
| $Q_{10}$ | Temperature coefficient for organic decay | Models 4b |
| $k_{ref}$ | Rate of decay at $T_{ref}$ | Models 4b |
| $T_{ref}$ | Reference temperature | Models 4b |
| *Wildlife Reservoir & Stress Dynamics* |  |  |
| $T_{cs}$ | Cold stress temperature threshold for wildlife breeding | Models 4a, 4b |
| $T_{hs}$ | Heat stress temperature threshold for wildlife breeding | Models 4a, 4b |
| $\beta_W$ | Population growth rate of wildlife reservoir | Models 4a, 4b |
| $\delta_W$ | Death rate of wildlife reservoir | Models 4a, 4b |
| $\sigma$ | Defecation rate of wildlife | Models 4a, 4b |
| $\alpha$ | Colonization rate from wildlife | Models 4a, 4b |
| $\chi$ | Scalar for drought stress effects on fungal dynamics | Model 4a, 4b |
| $S_{ds}$ | Soil moisture threshold for drought stress | Model 4a, 4b |
| $T_{ds}$ | Temperature threshold for drought stress | Model 4a, 4b |
| *Human Exposure & Epidemiology* |  |  |
| $\alpha_h$ | Human birth and immigration rate | Models 1–4b |
| $\omega$ | Human natural death and emigration rate | Models 1–4b |
| $c$ | Density-dependent human death rate | Models 1–4b |
| $\varepsilon$ | Infection rate parameter | Models 1–4b |
| $\psi, \psi_I$ | Rate of transition from exposed to symptomatic state | Models 2–4b |
| $\psi_A$ | Rate of transition from exposed to asymptomatic state | Model 4b |
| $\varrho, \varrho_I$ | Recovery rate from infection | Models 1–4b |
| $\varrho_A$ | Recovery rate from infection by asymptomatic | Model 4b |
| $\kappa$ | Disease-specific death rate | Models 1–4b |

| Parameter | Description | Models |
| --- | --- | --- |

We acknowledge that estimating fundamental biological traits such as optimal growth temperature and moisture from epidemiological data carries risks regarding parameter identifiability. However, this approach is necessitated by a critical gap in the empirical literature, as there is currently no high-resolution laboratory data sufficient to parameterize a continuous growth function for Coccidioides in soil. While historical studies like Friedman et al. [52] established broad survival limits, the data points are too sparse to derive a functional growth curve. Similarly, recent work by Mead et al. [53] provides growth rates at only two specific thermal set points (28°C and 37°C). Consequently, our fitted parameters (*T*_*opt*_, *S*_*opt*_) do not aim to replicate an in vitro fundamental niche, but rather quantify the pathogen’s realized niche in situ, the aggregate set of environmental conditions under which transmission effectively occurs in the complex, competitive soil matrix of the Sonoran Desert.

Regarding the use of scalars (*X*) and wildlife parameters (*β, T*_*cs*_), these are included not as ad-hoc fixes, but as mechanistic proxies for unmeasured but ecologically significant drivers. The drought stress scalar (*X*) tests the hypothesis that fungal proliferation is non-linearly amplified by extreme desiccation events, a dynamic that standard linear growth models fail to capture. Similarly, while we lack census data to validate the specific magnitude of the wildlife reservoir, fitting the wildlife parameters (*β*) allows the model to constrain the timing and shape of the fungal population curve based on host availability. By treating these as latent variables fit by human case data, we bridge the gap between soil ecology and spillover events, offering an efficient framework that accounts for pathogen amplification by animal reservoirs without requiring wildlife census data.

### 2.7 Statistical Baseline Model

To validate the predictive capability of our mechanistic approach, we compare the ODE models against the statistical framework established by Comrie [18] and Tamerius and Comrie [9]. This phenomenological approach assumes that *Coccidioides* abundance is driven by a two-phase climate sequence. First, there is a period of antecedent precipitation (typically 18 to 24 months prior), which enhances fungal saprobic growth in the soil, followed by concurrent arid conditions, high temperatures, and wind that facilitate spore aerosolization and dispersal. While Tamerius et al. [9] originally used negative binomial regression to account for overdispersion in count data, we performed a comparative analysis using both Generalized Linear Models (GLMs) with Poisson/Negative Binomial distributions and standard multivariate linear regression on incidence rates.

For this dataset, the GLM approaches exhibited convergence instability, higher Root Relative Mean Squared Error (RRMSE) in both fitting and forecasting, and higher Information Criteria than the linear approach. Consequently, we selected the multivariate linear regression model as the primary statistical baseline, as it provided the most accurate and stable error minimization. The baseline model forecasts monthly incidence rates (*I*_*t*_) based on concurrent temperature (*T*_*t*_) and *PM*_10_ (*P*_*t*_), alongside precipitation lagged by 18 and 24 months (*R*_*t*−18_, *R*_*t*−24_). The terms *β*_1_ and *β*_2_ are the regression coefficients for the concurrent drivers mean temperature (*T*_*t*_) and particulate matter concentration (*P*_*t*_, representing *PM*_10_), respectively. The coefficients *β*_3_ and *β*_4_ correspond to the drivers precipitation accumulation lagged by 18 months (*R*_*t*−18_) and 24 months (*R*_*t*−24_), while *ε*_*t*_ represents the residual error term. Thus, the model structure is

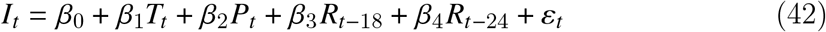

where predicted rates were constrained to be non-negative (*I*_*t*_ ≥ 0). This formulation allows for a direct comparison between biologically grounded mechanistic dynamics and standard meteorological correlation methods.

## 3 Methods

### 3.1 Data Sets

The Coccidioidomycosis human infected data is publicly available data of cases from Arizona, with all data available in the Supplementary Index. We selected the test regions of Pima County, Pinal County, Maricopa County, and the whole state of Arizona as four regions with slightly different population dynamics and a large enough number of Valley fever cases to merit modeling. The infected data we use is monthly, with the month associated with the case being the onset date entered into the Medical Electronic Disease Surveillance Intelligence System (MEDSIS), date reported to ADHS, or date submitted to the state. Monthly data was chosen to avoid the more stochastic nature of daily or weekly data, which can fluctuate in reporting due to events unrelated to the disease itself, like clinics and labs being closed for holidays. However, it is important to realize, fluctuations in the reported data may be artifacts of testing protocols and varying clinician awareness rather than true transmission dynamics.

The temperature and soil moisture data come from the National Oceanic and Atmospheric Administration (NOAA) climate at a glance webpage. The temperatures are the mean monthly temperatures for each specific county and the state as a whole, according to the NOAA data sources. The Palmer Z-Index was selected as the primary soil moisture biomarker because it effectively captures the short-term, upper-layer soil moisture anomalies that dictate the fungal saprobic cycle. By strictly balancing immediate precipitation with atmospheric evaporative demand, the Palmer Z-Index provides a highly biologically relevant proxy for soil moisture that drives Coccidioides hyphae growth and the subsequent arthroconidia [54].The Palmer Z-Index can be seen as a monthly moisture anomaly reflecting the deviation of current moisture conditions from normal conditions for a specified area. It is based on the principle of balancing moisture supply (precipitation and soil moisture) and moisture demand (potential evapotranspiration, recharge needed for soil, and runoff needed to maintain normal levels in streams, lakes, and reservoirs)[55]. The PM10 data sets come from the United States Environmental Protection Agency (EPA)[56], which is utilized exclusively for the statistical baseline model. Human population data was obtained from the 2010 and 2020 US Censuses, as well as the US Census Bureau’s estimates for 2011-2019 and 2021-2024.

### 3.2 Model Estimation

The models were built, run, and parameter fit in MATLAB 2025a. The large range for certain parameters, along with a large number of parameters, results in a vast search space for finding the optimal fit for the parameters. Basic algorithms within MATLAB like *fmincon* were unable to provide good fits for the data sets, with solutions getting caught around a local minimum. A more advanced algorithm, *particleswarm*, was utilized. The downside of *particleswarm* is that it requires intensive computing power in order complete a large number of runs in a reasonable amount of time. To improve convergence and computational efficiency, the *particleswarm* optimization was parallelized and coupled with a hybrid function (fmincon) for final parameter refinement. Additionally, because the environmental and epidemiological dynamics can cause numerical stiffness and widely varying computation times, the evaluation dynamically shifted between non-stiff (ode45, ode78) and stiff (ode15s, ode23s) solvers to prevent stalling.

### 3.3 Model Fitting Comparison

First, to quantify the goodness-of-fit for each model, we utilize the Root Relative Mean Square Error (RRMSE), which is calculated as

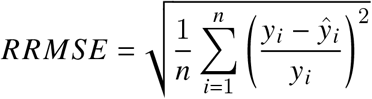

where *n* is the number of observations, *y*_*i*_ is the observed case data at time *i*, and *ŷ*_*i*_ is the model-predicted value. RRMSE was chosen over the more standard Root Mean Square Error (RMSE) because we assume epidemiological data, like the Valley fever cases used here, is fundamentally heteroscedastic. Heteroscedasticity means the variance of the error (the noise) is not constant and tends to increase as the number of infected cases increases. With RRMSE, by normalizing the error as a percentage of the observed value, provides a more balanced assessment of model performance across both low-case troughs and high-case peaks [57, 58]. This approach is analogous to a weighted least squares, where each squared error is weighted by 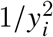. The assumption that the relative errors are approximately normally distributed can be assessed using Relative Quantile-Quantile (QQ) plots [59], which are included in the Supplementary Index. Under this formulation, the corresponding errors on the original scale are shown to exhibit heteroscedasticity.

Simply choosing the model that best fits the data (the one with the lowest RRMSE) is insufficient for making public health decisions, as more complex models will almost always fit the training data better, but may be overfitting the noise [60]. This trade-off between model fit (low bias) and model complexity (low variance) is where formal model selection tools in the form of Akaike Information Criterion (AIC), Akaike Information Criterion corrected (AICc), and Bayesian information criterion (BIC) become excellent tools for the job [61]. AIC, AICc, and BIC are penalized-likelihood criteria. All three start with a measure of how well the model fits the data, the likelihood, and then add a penalty term that increases with the number of parameters in the model. We are assuming the epidemiological data to be heteroscedastic, so calculating the likelihood via the standard Residual Sum of Squares (RSS) would violate our error variance assumptions and fail to yield the true Maximum Likelihood Estimate for this system. Instead, the Information Criteria were computed directly from the weighted residuals. To ensure consistency between the estimation procedure and model selection criteria, we assume that the relative errors are independent and approximately normally distributed. Under this assumption, maximizing the likelihood is equivalent to minimizing the sum of squared relative errors, which corresponds directly to minimizing the squared RRMSE. The log-likelihood was therefore constructed from the weighted residuals induced by this relative error formulation, ensuring that the AIC, AICc, and BIC are computed consistently with the estimation objective. This formulation ensures that the model selection criteria are evaluated consistently with the parameter estimation procedure, while incorporating a heteroscedastic variance structure induced by the relative error model.

Akaike Information Criterion (AIC) seeks to optimize model selection by minimizing the Kullback-Leibler divergence, which measures the information lost when a model is used to approximate the true data-generating process [62, 61]. The corrected version, AICc, is especially advantageous for smaller sample sizes, where the number of data points *n* is not large relative to the number of parameters *K*, as it imposes a greater penalty for additional parameters, thus mitigating the risk of overfitting [61]. Bayesian Information Criterion (BIC), based on Bayesian inference, is formulated to favor simpler models (those with fewer parameters) compared to AIC, particularly in the context of larger sample sizes [63]. Its objective is to identify the model that is most probabilistically aligned with the true underlying data-generating process among the set of competing models. In model selection, the model exhibiting the lowest values of AIC, AICC, or BIC is deemed the most preferable [61].

### 3.4 Forecasting

We utilize expanding window forecasting [64, 65] which mimics a realistic forecast scenario. An expanding window forecast for our purposes means, for our data set of eleven years of infected humans, we fit the first eight years (January 2013-January 2021) of data and forecast the ninth year (January 2021-January 2022), then fit the first through the ninth year (January 2013-January 2022) and forecast the tenth year (January 2022-January 2023), and finally fit the first through the tenth year (January 2013-January 2023) and forecast the eleventh year (January 2023-January 2024). We calculate the RRMSE of the forecasted ninth, tenth and eleventh year and the average RRMSE across the years. This method is a scaled back version of time-series cross-validation laid out by Hyndman and Athanasopoulos [64].

### 3.5 Forecasting Accuracy Testing

To statistically compare the predictive accuracy of the two models, we first utilized the Diebold-Mariano (DM) test [66]. While RRMSE was strictly necessary during parameter estimation to account for the heteroscedasticity of the long-term training data, forecasting evaluation requires a different clinical perspective. In public health, a large relative error during a low-incidence disease trough is epidemiologically less consequential than a large absolute error during a severe seasonal outbreak. Therefore, to align our model validation with actual public health resource allocation priorities, we evaluate the out-of-sample forecasts using standard absolute squared loss differentials. Let *e*_*Stat,t*_ and *e*_*ODE,t*_ denote the raw forecast errors (*y*_*t*_ − *ŷ*_*t*_) for the Statistical Baseline Model and the ODE model (Models 1–4b), respectively, at time *t*. To compute the test statistic, we calculate the loss differential at each time point *t* as 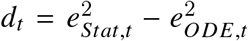. The test assesses the sample mean of these differentials, 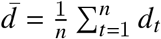, standardized by a consistent estimate of its long-run variance, 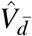, which accounts for serial correlation up to the forecast horizon *h* − 1:

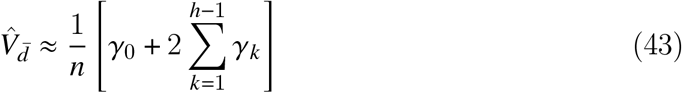

where *γ*_*k*_ is the autocovariance of *d*_*t*_ at lag *k*. The standard Diebold-Mariano test statistic is then defined as 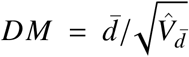, which under the null hypothesis follows a standard normal distribution asymptotically. Formally, we use this statistic to evaluate the null hypothesis of equal predictive accuracy against the two-sided alternative. The hypotheses are defined as follows:

- **Null Hypothesis (***H*_0_): The two models exhibit equal predictive accuracy.

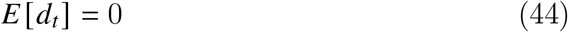
- **Alternative Hypothesis (***H*_1_): The two models exhibit different levels of predictive accuracy.

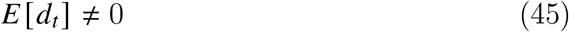

If the null hypothesis is rejected, we conclude that the difference in forecast accuracy between the Statistical Baseline and the ODE model is statistically significant.

Although the standard DM test statistic assumes asymptotic normality, it is known to be oversized (rejecting the null hypothesis too frequently) in samples of moderate size, particularly when the forecast horizon, *h*, is greater than 1. Given that our analysis involves a forecast horizon of *h* = 12 months, the forecast errors exhibit strong serial correlation. To address this and mitigate finite-sample bias, we also employed the Modified Diebold-Mariano (MDM) test [67]. The MDM test adjusts the DM test statistic using a correction factor that accounts for the sample size, *n*, and the forecast horizon, *h*:

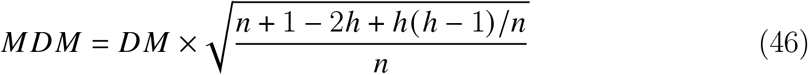

By incorporating this correction factor, the MDM statistic penalizes the test for longer forecast horizons and smaller sample sizes. Furthermore, rather than comparing the test statistic to the standard normal distribution, the MDM statistic is compared to the critical values of the Student’s *t*-distribution with *n* − 1 degrees of freedom, providing a more conservative and robust inference for our forecasting evaluation.

### 3.6 Sensitivity Analysis

To evaluate the robustness of each model and identify the parameters that exert the most influence on the respective dynamics of each system, a comprehensive sensitivity analysis was conducted. Sensitivity analysis is crucial for understanding which factors are primary drivers of disease outcomes and where efforts to refine parameter estimates would be most impactful [68, 69]. We utilize the method of Integrated Absolute Normalized Sensitivity (IANS), which was chosen for its robustness in quantifying the total, accumulated impact of a parameter’s variation over the entire simulation period [70]. The IANS for a specific state variable, *x*_*i*_ (such as *I*), with respect to a parameter, *p*_*j*_ (such as *ω*), is calculated by integrating the absolute value of the normalized sensitivity coefficient over time:

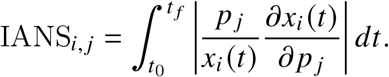

To compute the partial derivatives 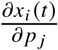 continuously over time, we utilized Forward Sensitivity Analysis (FSA). The sensitivity equations were dynamically formulated and integrated simultaneously with the primary state equations using MATLAB’s internal ODE sensitivity framework, avoiding the truncation errors inherent in finite difference approximations. A large IANS value signifies that the parameter *p*_*j*_ has a strong and sustained influence on the state *x*_*i*_, making IANS a strong metric for assessing overall parameter importance in complex systems [71]. For this analysis, we focus specifically on the sensitivity of the Infected Humans (*I*) compartment. Infected Humans are the primary public health interest and represent the observable data that the models aim to fit and forecast. Thus, understanding which parameters most significantly drive the dynamics of the infected population is the goal of this sensitivity analysis.

## 4 Results

We present the results of the model fittings, forecasting results, and sensitivity analysis.

### 4.1 Model Fitting

The RRMSE model-fitting results are presented for the four regions (Arizona, Maricopa County, Pima County, Pinal County) in Figures 5-8. These figures and the summary in Table 2 indicate a clear trend in which the RRMSE score improves as mechanistic complexity increases. The fitting figures visually corroborate this RRMSE quantitative improvement. The following figures compare the fitted model outputs, represented with solid lines, against the observed monthly Coccidioidomycosis cases, shown as dots for each of the four study regions.

**Table 2:** The RRMSE of the fit for each model for each region and the mean RRMSE across all regions.

| Model | Arizona | Maricopa County | Pima County | Pinal County | Mean |
| --- | --- | --- | --- | --- | --- |
| Model 1 | 2.2534% | 2.3731% | 2.1144% | 2.8401% | 2.3953% |
| Model 2 | 2.2475% | 2.3679% | 1.9781% | 2.8375% | 2.3578% |
| Model 3 | 1.5679% | 1.3506% | 1.7256% | 2.6570% | 1.8253% |
| Model 4a | 1.5133% | 1.3606% | 1.5501% | 1.5784% | 1.5006% |
| Model 4b | 1.3107% | 1.2547% | 1.4724% | 1.5675% | 1.4013% |
| Statistical | 2.6701% | 2.6865% | 2.2811% | 3.1097% | 2.6869% |

**Figure 5:**
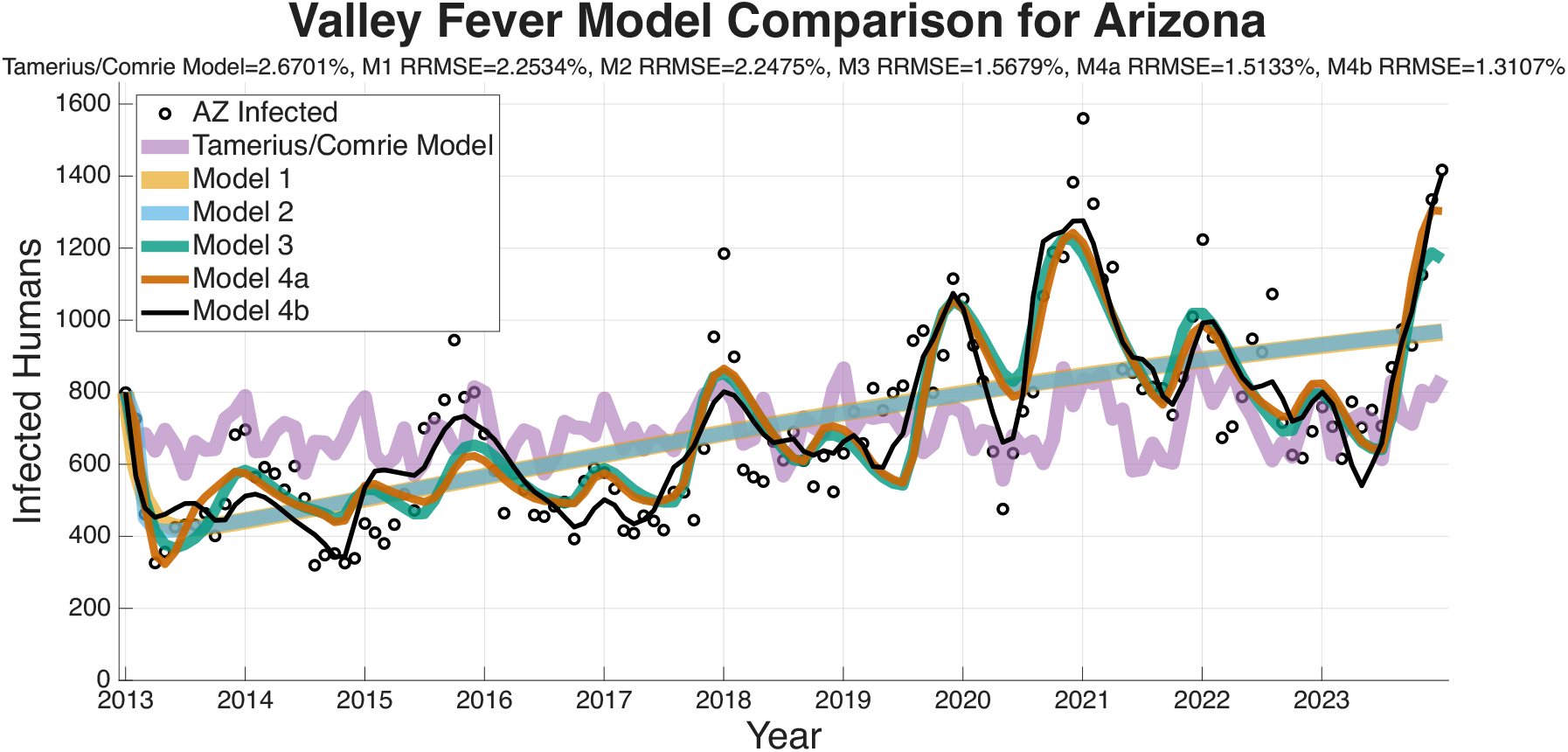
Comparison of the five fit ODE models and Tamerius/Comrie Statistical Baseline model against observed infected human data for the entire state of Arizona. The observed cases are shown as small black circles, while each model fit is a colored line. Models 4a (brown/orange line) and 4b (black line) demonstrate the closest visual fit to the seasonal peaks and valleys of the data.

In Figure 5, the fits for the entire state of Arizona are shown. Model 1 (brown/orange) and Model 2 (light blue) only capture the broad, long-term trend, failing to replicate the seasonal variability. Model 3 (turquoise) introduces seasonality with the inclusion of climate effect functions, but consistently misses the magnitude of the peaks and valleys. Models 4a and 4b provide a much closer fit to the intra-annual fluctuations. Model 4b achieved the lowest RRMSE for this region at 1.3107%.

Figure 6 displays the model fits for Maricopa County, which has the highest number of cases of any county in Arizona and the largest population. The Pima County data in Figure 7 is interesting as Model 3 (turquoise) has a notably poor fit, particularly in the early years of the dataset, resulting in a RRMSE (2.0081%). This suggests that the environmental drivers in Model 3, without the modulation from the wildlife compartment, are insufficient to capture Pima County’s specific dynamics. Models 4a (1.5501% RRMSE) and 4b (1.4724% RRMSE) correct this, suggesting that including the wildlife component substantially improves the model’s ability to capture the observed dynamics. Figure 8 shows the fits for Pinal County, the least populous county we fit the models to, and a region with sporadic case peaks. As with the other regions, Models 1 and 2 only capture the basic upward trend. Models 3, 4a, and 4b all effectively capture the seasonal nature of the data. However, Models 4a and 4b, with RRMSEs of 1.5784% and 1.5675%, respectively, are more accurately able to match the timing and magnitude of the peaks post-2019. Across all regions, the Tamerius/Comrie Statistical Baseline model (light purple) has peaks and valleys at some of the right times but does not ever go low or high enough to capture the extreme dynamics of the disease. This visual evidence supports the quantitative RRMSE scores, where Model 4b is again selected as the best fit.

**Figure 6:**
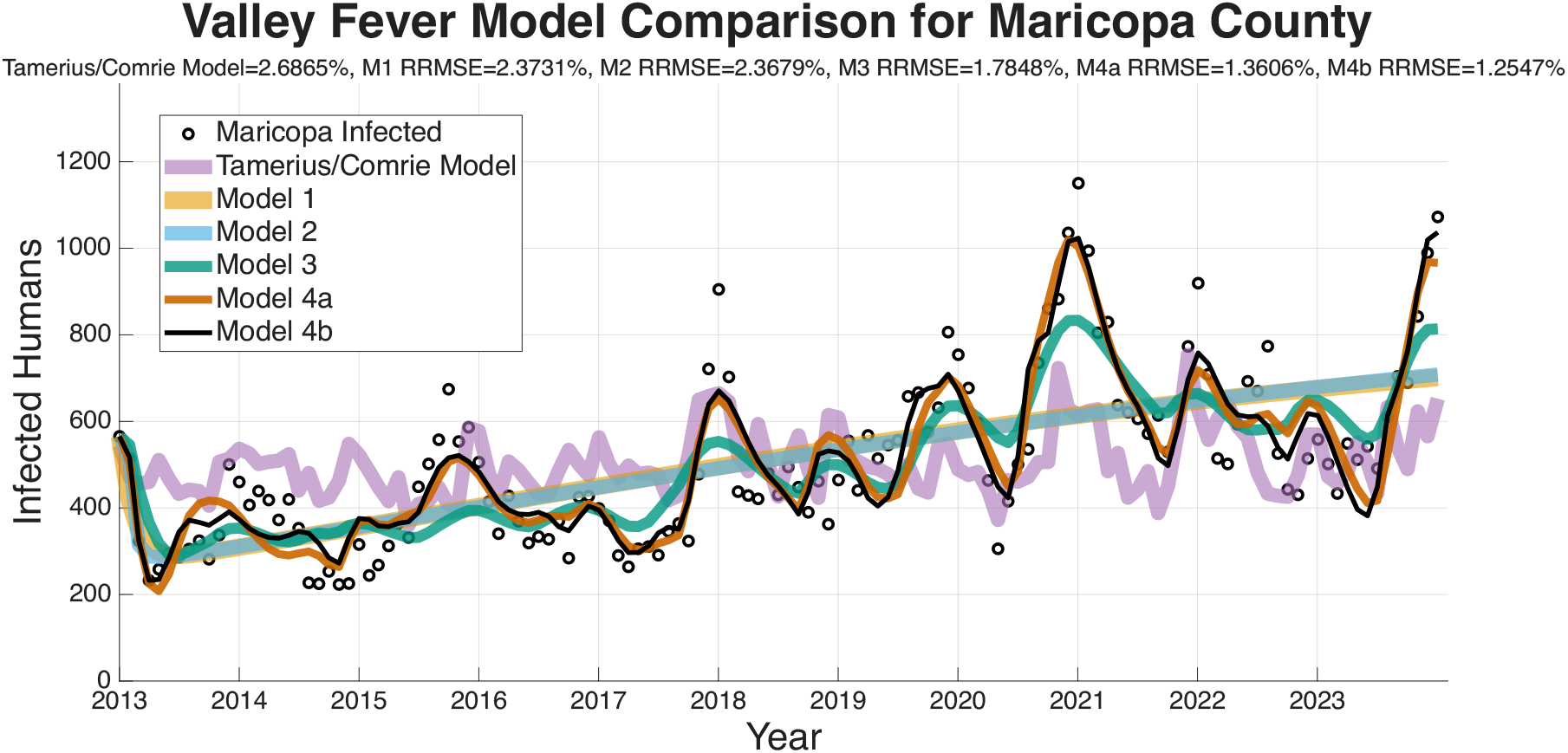
ODE and Statistical model fit comparison for Maricopa County, the highest population county in Arizona.

**Figure 7:**
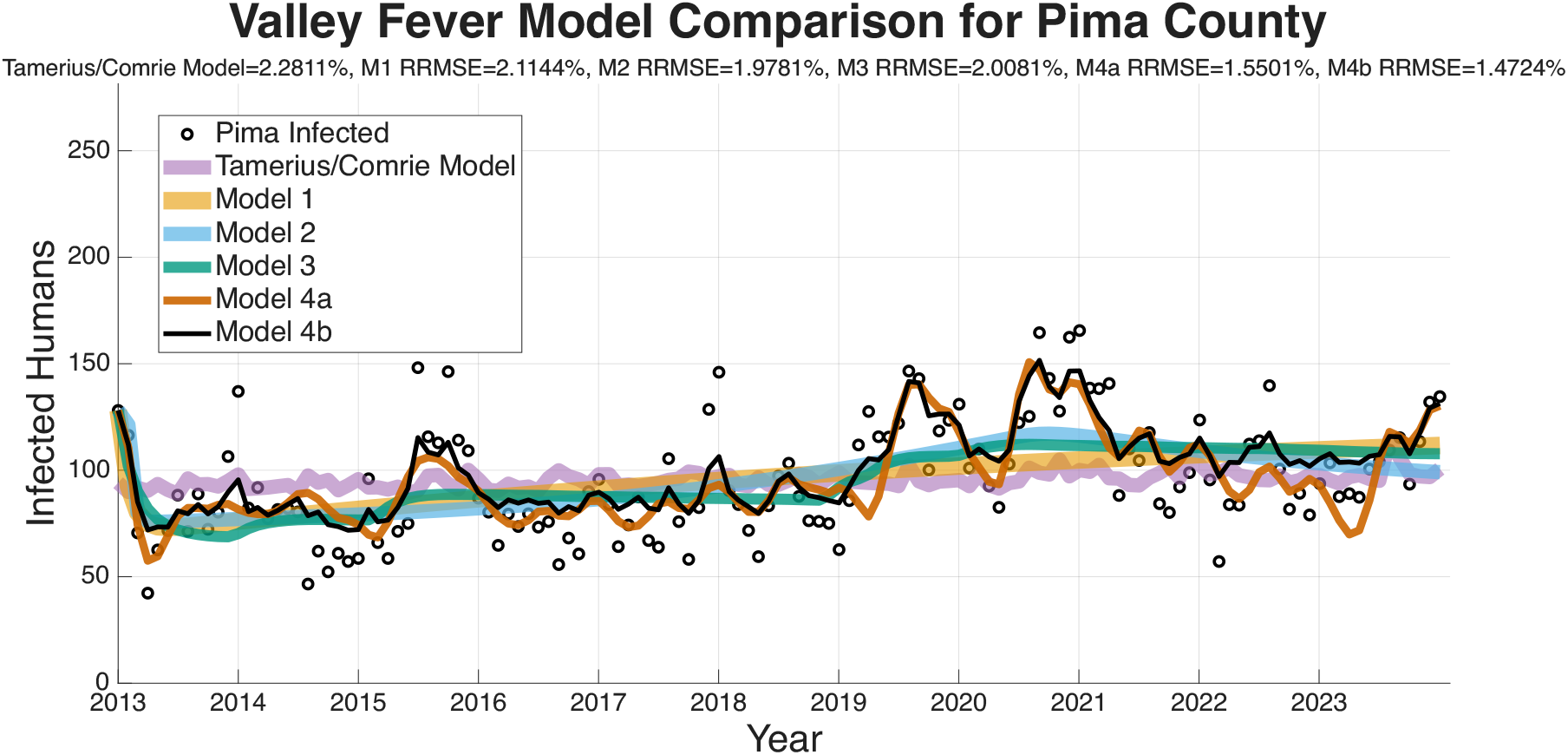
ODE and Statistical model fit comparison for Pima County.

**Figure 8:**
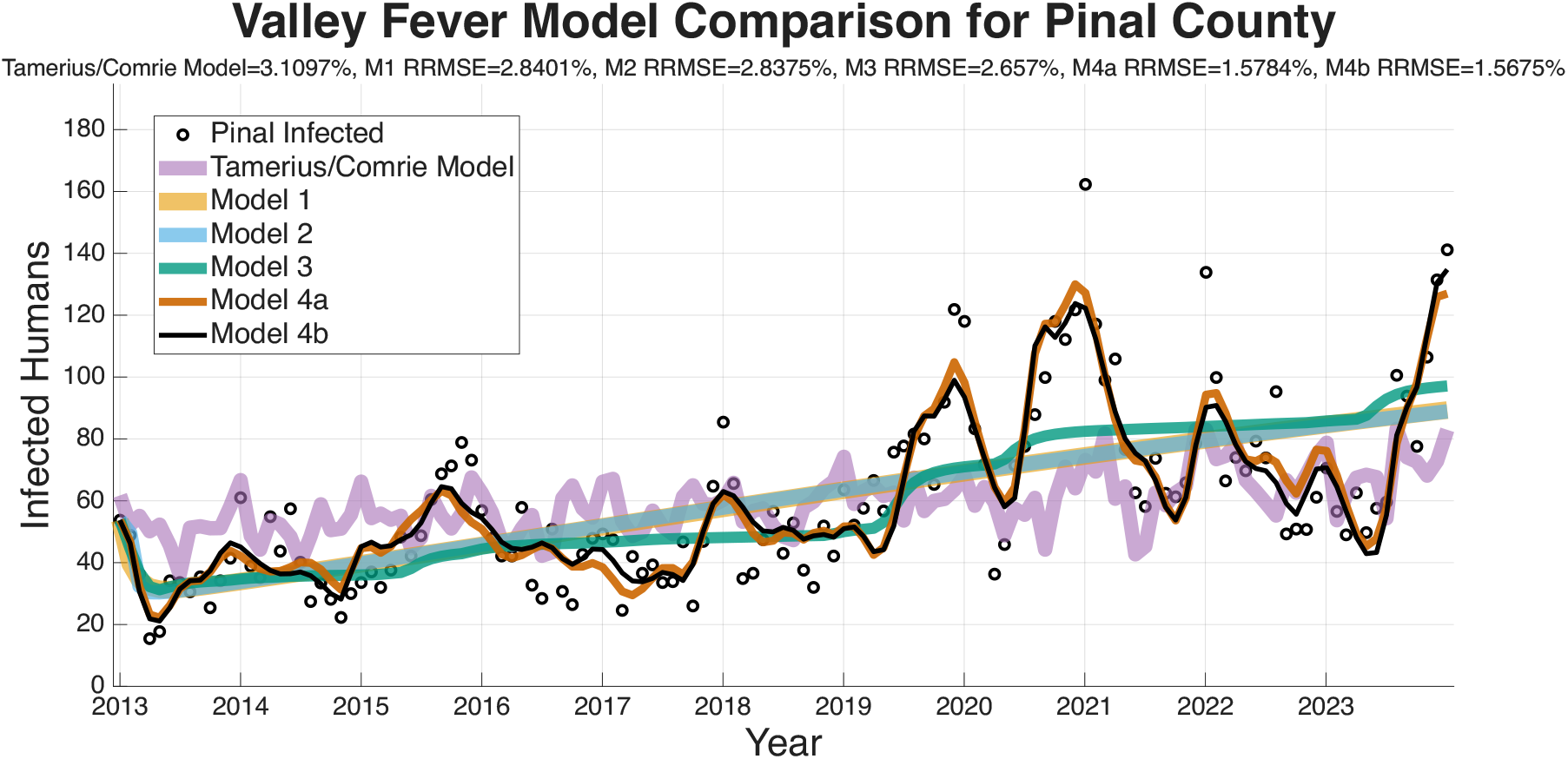
ODE and Statistical model fit comparison for Pinal County, the least populated of the counties we are utilizing.

In summary, a consistent visual pattern emerges across all four geographic regions. Models 1 and 2 are only able to fit the rough, multi-year mean trend and fail to capture the critical intra-annual variability. Models 3, 4a, and 4b demonstrate a qualitatively superior ability to capture this within-year seasonality. While Model 3 is a significant improvement, it performs relatively poorly compared to the more complex models at fitting the specific peaks and valleys in the observed case data. Visually, Models 4a and 4b show very similar and tight fits to the data, confirming the RRMSE scores that select them as the most effective models.

This conclusion is robustly supported by the information criterion scores, which balance goodness-of-fit with model simplicity as the inclusion of additional parameters are penalized. Across the three Information Criteria, Akaike Information Criterion (AIC), corrected Akaike Information Criterion (AICc), and Bayesian Information Criterion (BIC), Model 4b consistently yields the most preferable (lowest) mean score (Table 3). This shows that the superior fit of Model 4b is supported even after accounting for the complexity of the model. Since the improvement is not solely attributable to overfitting, the increased complexity represents a statistically justified enhancement in model structure. Notably, the most significant drop in criterion scores occurs between Model 2 and Model 3, underscoring the critical importance of incorporating environmental drivers (*F*_*H*_ (*T, S*_*m*_) and *F*_*A*_ (*T, S*_*m*_)) to adequately explain the system’s dynamics.

**Table 3:**
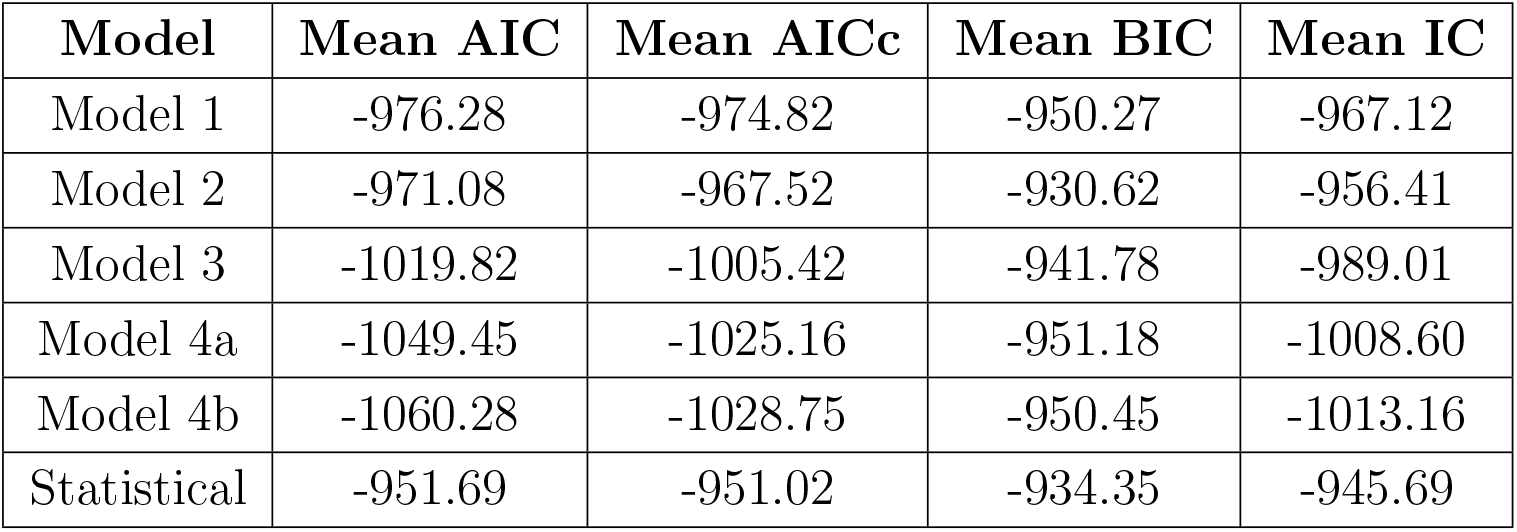
The columns labeled Mean AIC, Mean AICc, and Mean BIC show the mean information criterion score of the respective type across Maricopa County, Pima County, Pinal County, and the state of Arizona. The final column labeled Mean IC is the mean of the mean AIC, mean AICc, and mean BIC. Note that highly negative values indicate a stronger model fit relative to complexity.

Because the likelihood estimates are derived from the Root Relative Mean Square Error (RRMSE), the resulting information criteria scores (AIC, AICc, and BIC) are negative. They are negative because the RRMSE values represent relative, proportional errors that are strictly between 0 and 1. For example, an RRMSE of 1.5% is evaluated as 0.015. The model selection criteria rely on the natural logarithm of the estimated relative variance, ln(RRMSE^2^), and the natural logarithm of any value between 0 and 1 is inherently negative. As a model’s fit improves and its error variance approaches zero, the logarithmic term approaches negative infinity, resulting in a more negative base score. The penalty for model complexity adds a positive value, pulling the score upward. Therefore, the standard rule that models with lower information criterion values are preferred remains unchanged, with comparisons interpreted relative to competing models. The model that has the optimal balance between minimizing predictive error and avoiding over-parameterization is the one that yields the lowest or most negative overall score.

### 4.2 Forecasting Human Cases

The ability of a mathematical model to accurately forecast future dynamics is a central objective in epidemiology and many applied modeling contexts. As such, we test the ability of the models we have outlined to see which are capable of producing reliable forecasts. The following figures for the four regions we are forecasting (Arizona in Figure 9, Maricopa county in Figure 10, Pima county in Figure 11, and Pinal county in Figure12) show off the forecasts for each expanding window period. In each figure (9–12) January 2013-December 2020 is fit, January 2021-December 2021 is forecasted, January 2013-December 2021 is fit, January 2022-December 2022 is forecasted, January 2013-December 2022 is fit, January 2023-December 2023 is forecasted.

**Figure 9:**
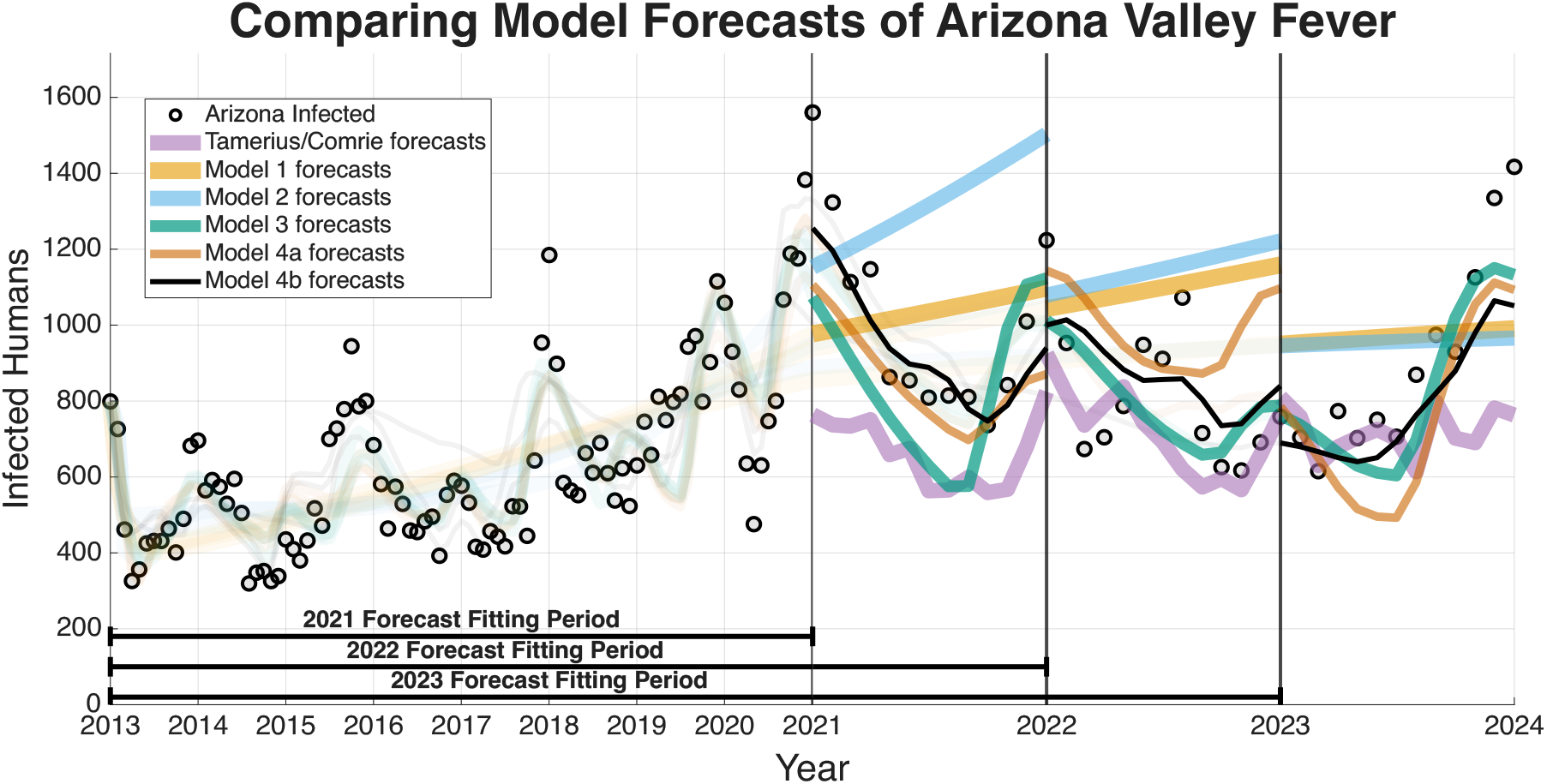
Arizona model forecasting results across all models.

**Figure 10:**
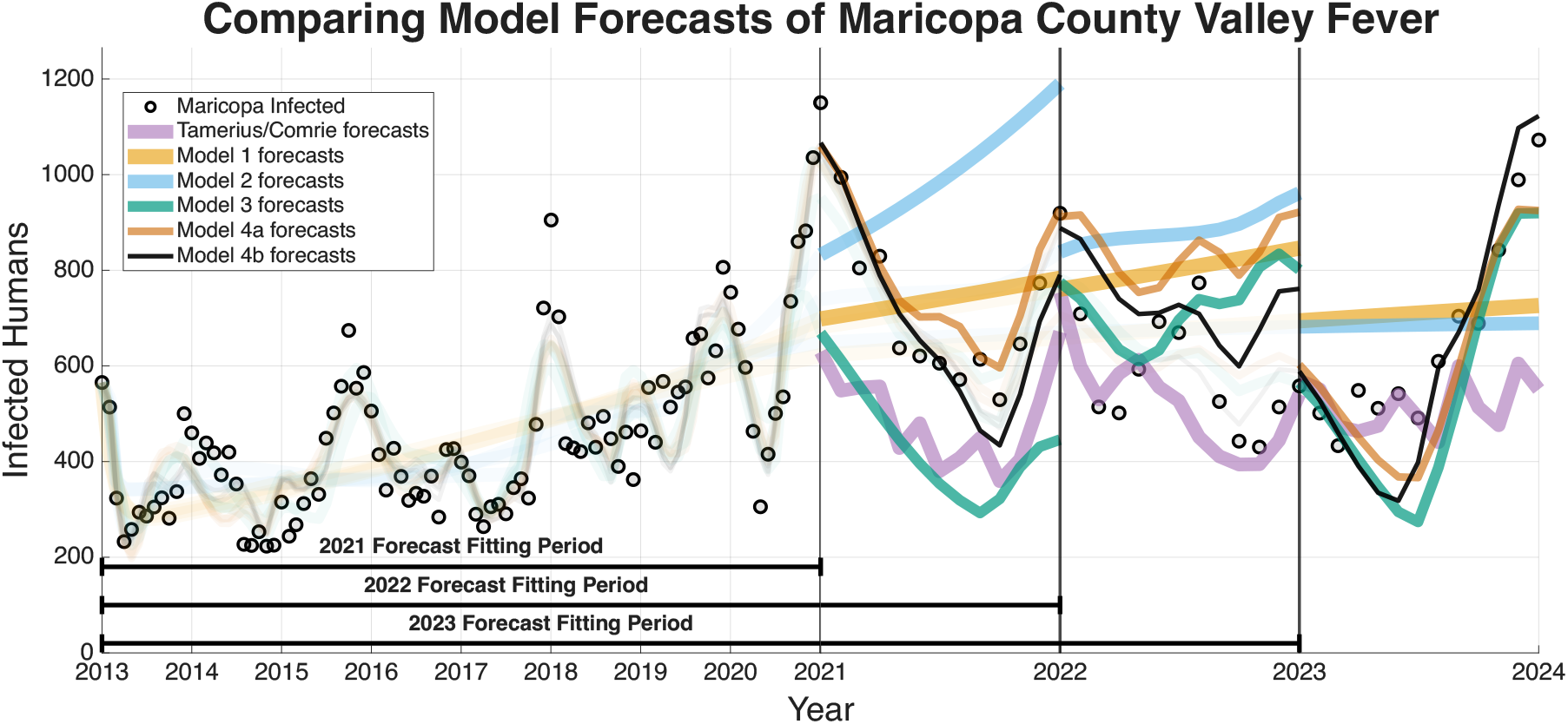
Maricopa county model forecasting results across all models.

**Figure 11:**
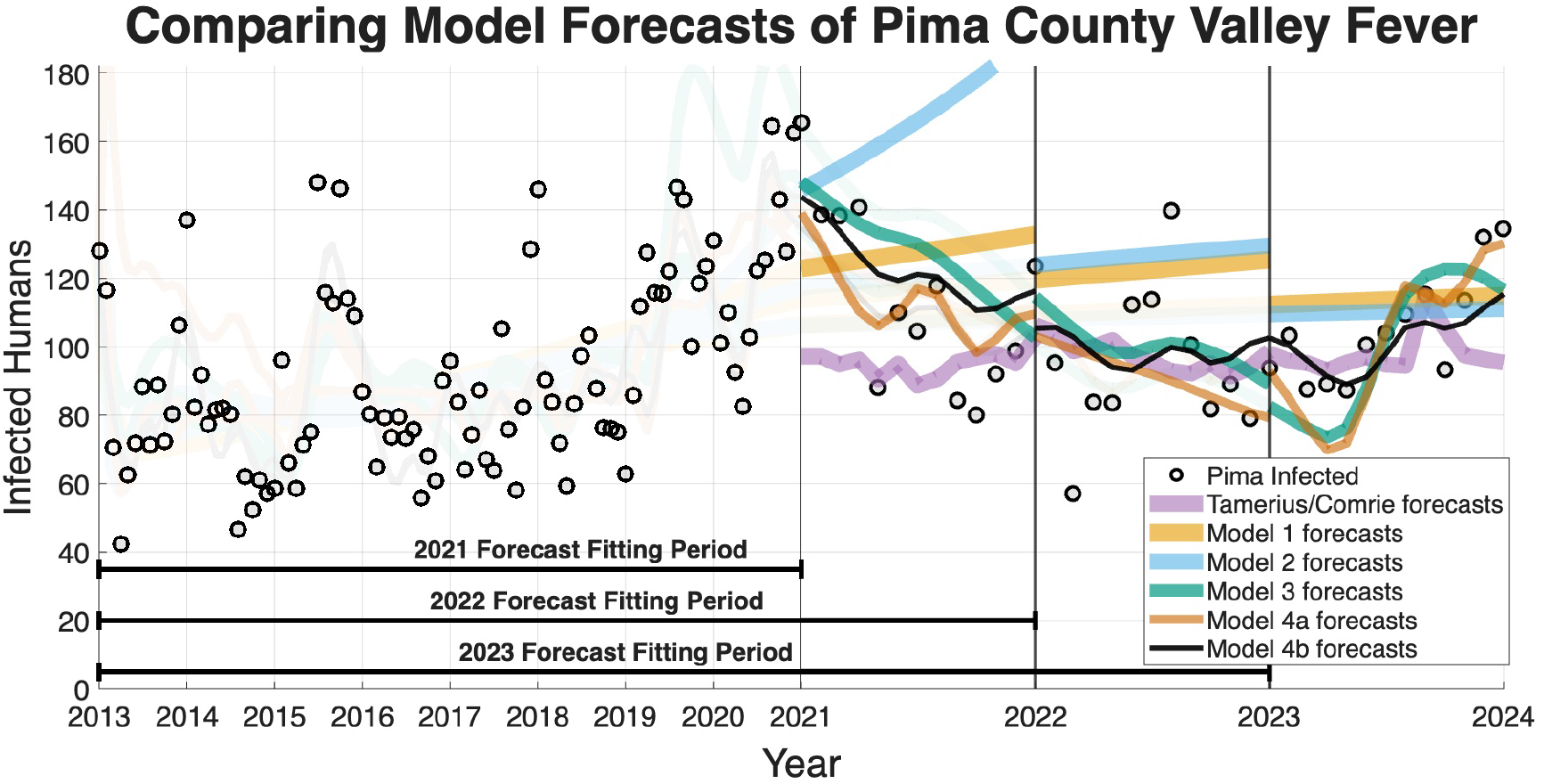
Pima county model forecasting results across all models.

Figures 9–12 visually suggest that the mechanistic models, particularly those incorporating wildlife reservoir dynamics (Models 4a and 4b), more closely track the timing and magnitude of seasonal outbreaks. However, we need to quantify this predictive performance. Table 4 aggregates the results, presenting the mean forecasted RRMSE across the three expanding windows forecasted years (2021, 2022, and 2023) for each model and geographic region. The full year-by-year RRMSE forecasting results for each specific expanding temporal window are available in the Supplementary Information.

**Figure 12:**
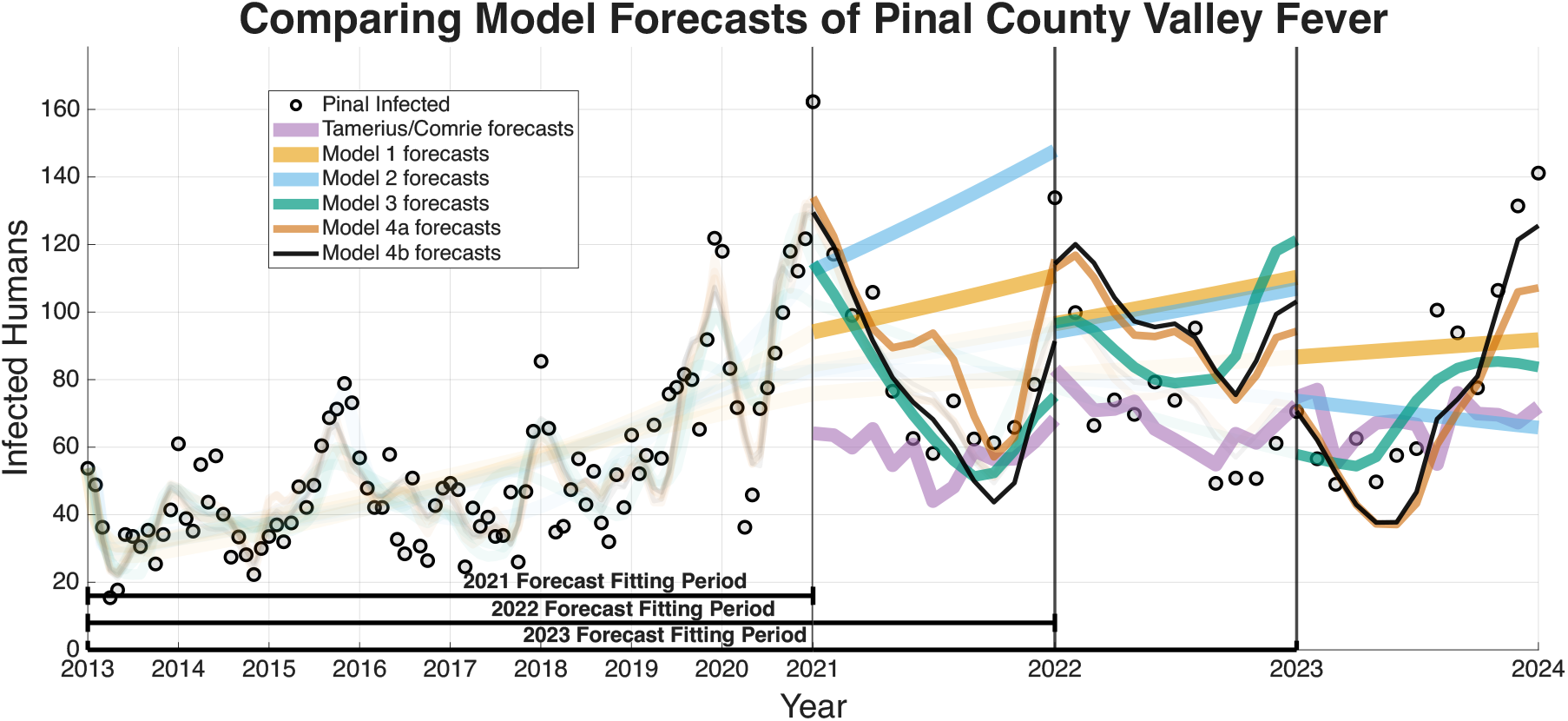
Pinal county model forecasting results across all models.

**Table 4:**
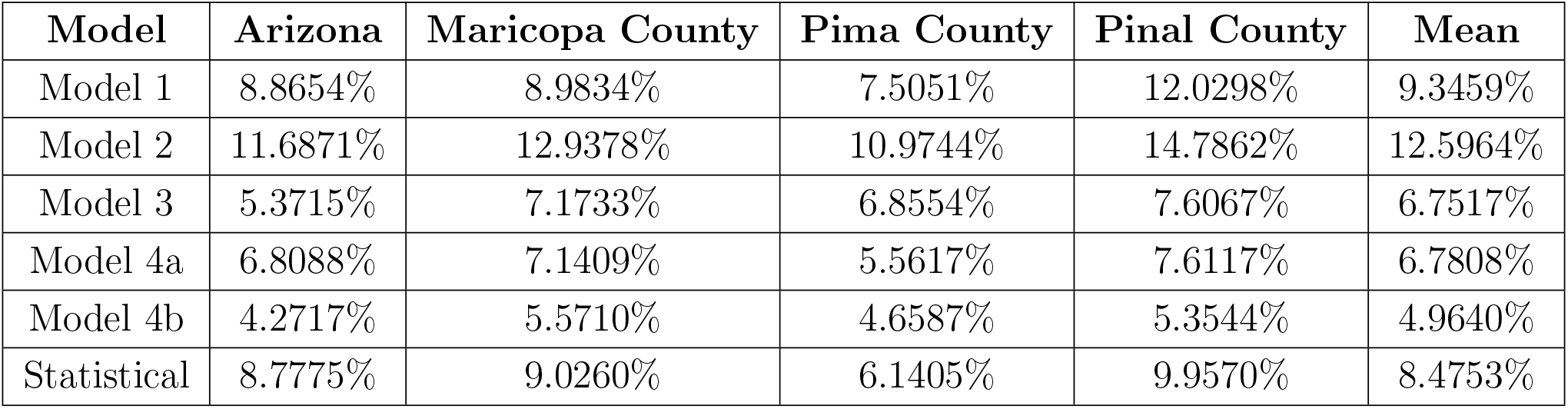
Mean RRMSE across the three expanding window forecasted years of 2021, 2022, and 2023.

| **Model** | **Arizona** | **Maricopa County** | **Pima County** | **Pinal County** | **Mean** |
| --- | --- | --- | --- | --- | --- |
| Model 1 | 8.8654% | 8.9834% | 7.5051% | 12.0298% | 9.3459% |
| Model 2 | 11.6871% | 12.9378% | 10.9744% | 14.7862% | 12.5964% |
| Model 3 | 5.3715% | 7.1733% | 6.8554% | 7.6067% | 6.7517% |
| Model 4a | 6.8088% | 7.1409% | 5.5617% | 7.6117% | 6.7808% |
| Model 4b | 4.2717% | 5.5710% | 4.6587% | 5.3544% | 4.9640% |
| Statistical | 8.7775% | 9.0260% | 6.1405% | 9.9570% | 8.4753% |

We also recognize that a lower aggregated RRMSE alone does not guarantee a scientifically meaningful improvement in real-world forecasting accuracy. Thus, we formalize our comparative evaluation using standard absolute loss differentials in the form of Diebold-Mariano (DM) and Modified Diebold-Mariano (MDM) tests. To increase statistical power, forecast errors were pooled across the four regions, treating each region’s month as an observation in the loss differential series. While state-level data partially aggregates county-level dynamics and thereby introduces some dependence between observations, the regions are modeled independently and exhibit distinct patterns. As such, this pooled analysis provides an aggregate assessment of predictive performance across multiple spatial scales, though the resulting inference should be interpreted as a combined rather than strictly independent comparison. Table 5 and Table 6 outline the results of the DM and MDM tests, respectively, which systemically assess whether the predictive superiority of the ODE frameworks over the phenomenological baseline is statistically significant.

**Table 5:** Diebold-Mariano (DM) test comparing the Baseline Statistical Model to each of the ODE models across three years of forecasting for the four regions (N=144).

| **Model 1** | **Model 2** | **Model 3** | **Model 4a** | **Model 4b** |
| --- | --- | --- | --- | --- |
| DM= -0.9312 | DM=-3.9891 | DM =1.2608 | DM =2.1809 | DM = 3.2101 |
| p=0.3517 | p=0.0001 | p=0.2074 | p=0.0394 | p=0.0001 |
| No Sig. Difference | Baseline is better | No Sig. Difference | ODE is better | ODE is better |

**Table 6:** Modified Diebold-Mariano (MDM) test comparing the Baseline Statistical Model to each of the ODE models across three years of forecasting for the four regions (N=144).

| **Model 1** | **Model 2** | **Model 3** | **Model 4a** | **Model 4b** |
| --- | --- | --- | --- | --- |
| MDM = -0.4908 | MDM = -2.0491 | MDM = 0.8191 | MDM =1.2353 | MDM =1.8613 |
| p = 0.6243 | p = 0.0423 | p = 0.4141 | p =0.0264 | p =0.0012 |
| No Sig. Difference | Baseline is better | No Sig. Difference | ODE is better | ODE is better |

The Information Criteria established that Model 4b provides the most statistically justified fit to the training data, outperforming the Statistical Baseline Model across all metrics, including AIC, AICc, and BIC. To confirm that this structural superiority translates to true predictive power, we evaluated the out-of-sample expanding window forecasts. To determine if the visual forecasting improvements over the Statistical Baseline were statistically significant, we conducted a Diebold-Mariano (DM) test and a Modified Diebold-Mariano (MDM) test as well. The MDM test is particularly robust for our purposes, as our forecasting horizon extends up to 12 months. This test adjusts for the inherent serial correlation of forecast errors, particularly for longer forecast horizons. The tests yielded statistics and corresponding *p*-values as seen in Tables 5 and 6. For Models 4a and 4b, the tests resulted in positive test statistics with *p* < 0.05, leading us to reject the null hypothesis of equal predictive accuracy. This statistically confirms that the inclusion of mechanistic ecological and wildlife reservoir drivers in these advanced models significantly outperforms the phenomenological Statistical Baseline in terms of absolute forecasting accuracy, particularly during critical, high-incidence seasonal outbreaks.

### 4.3 Sensitivity Analysis

For the sensitivity analysis we are calculating the Integrated Absolute Normalized Sensitivity (IANS), a dimensionless value, for each parameter in the respective models in respect to the infected human compartment. The results of the IANS Sensitivity analysis show that population parameters *α*_*h*_ (birth/immigration) and *ω* (death/migration) exhibit the greatest influence on model output for Models 1–4a, and are highly sensitive for Model 4b, as seen in Figures 13-17. This is not unusual, as general population dynamics can be the strongest driver of any diseased subpopulation with a low death rate over a large enough time span.

**Figure 13:**
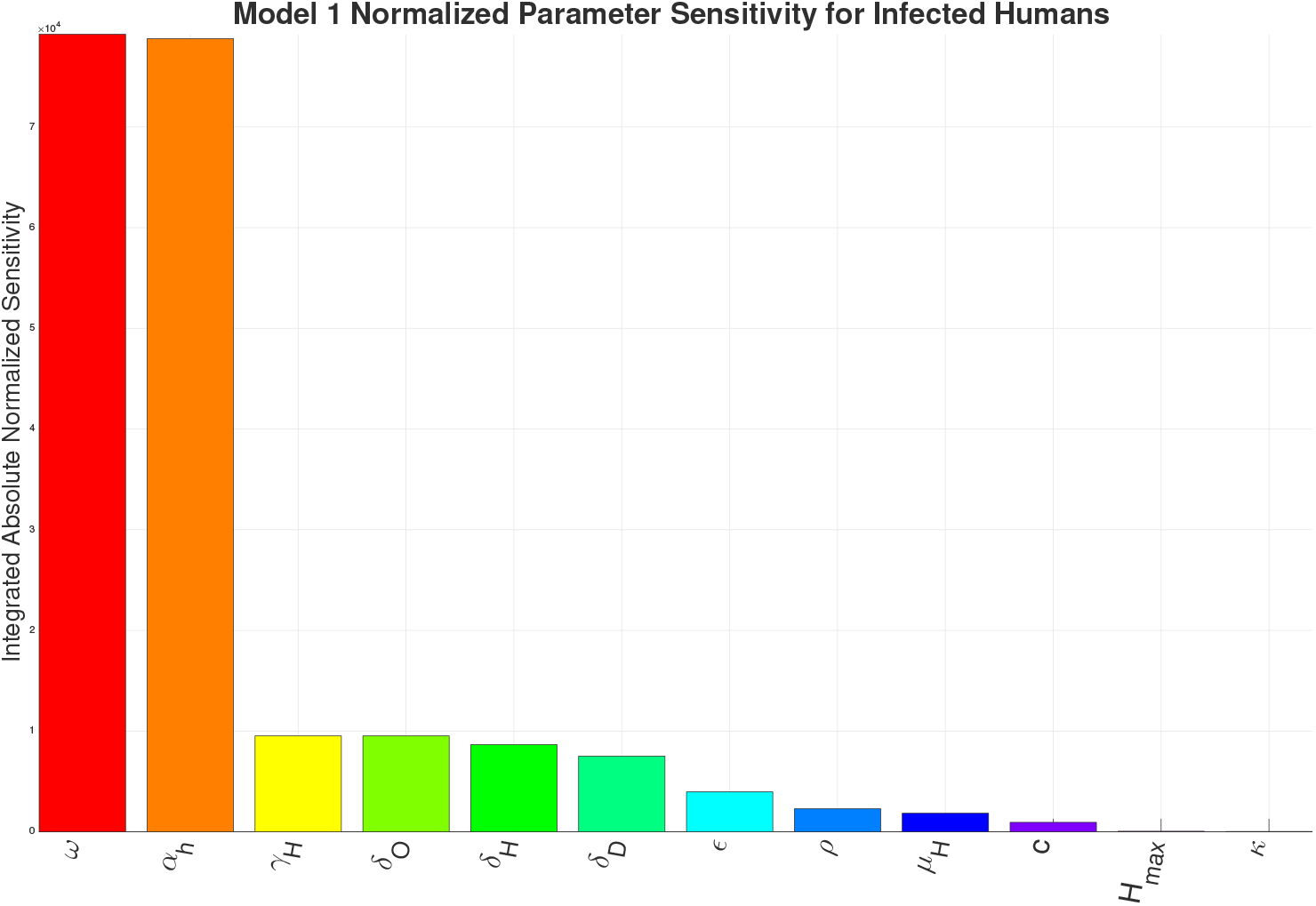
Model 1 Integrated Absolute Normalized Sensitivity analysis.

In Figure 13 (Model 1) and Figure 14 (Model 2), *γ*_*H*_ the growth rate of hyphae, is the most sensitive parameter behind the population parameters. With the introduction of the environment in Model 3, environmental parameters such as 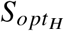 (optimal soil moisture level for hyphae), 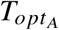 (optimal temperature for arthroconidia), 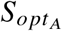 (optimal soil moisture level for arthroconidia), and 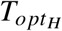 (optimal temperature for hyphae) as seen in Figure 15, become the most sensitive behind the population parameters. For Model 4a and Model 4b, as seen in Figure 16 and Figure 17, which introduce the wildlife interaction with *Coccidioides*, the wildlife parameters of *T*_*hs*_ and *T*_*cs*_ become the most highly sensitive non-human population parameters, even to the point of becoming the most sensitive parameters overall in Model 4b. These parameters control the population growth time of wildlife. The parameters *T*_*hs*_ and *T*_*cs*_ are the most sensitive in Model 4b, highlighting the influence of the wildlife population growth function and suggesting that including robust wildlife data and better empirical characterization of wildlife dynamics could improve model fidelity.

**Figure 14:**
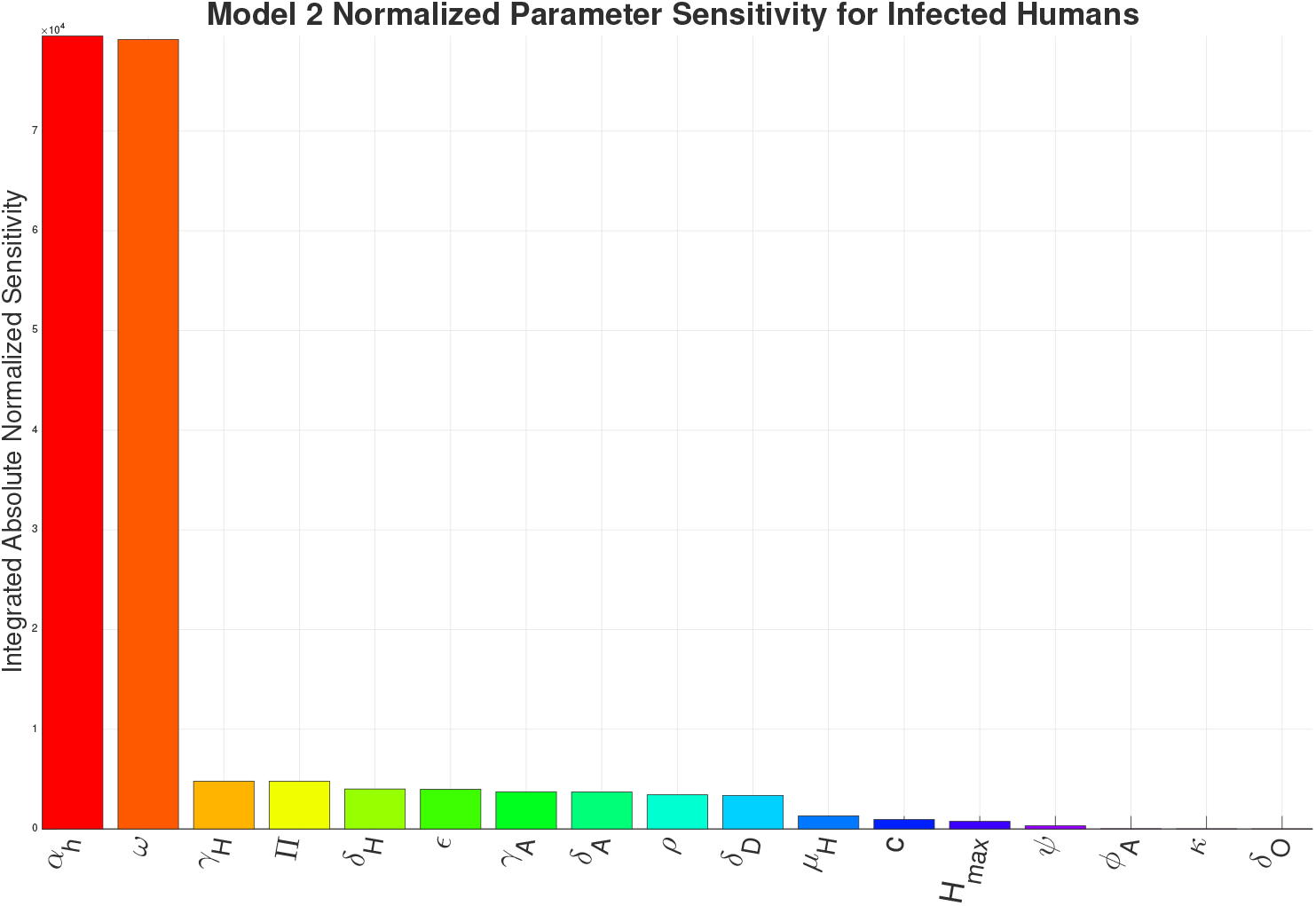
Model 2 Integrated Absolute Normalized Sensitivity analysis.

**Figure 15:**
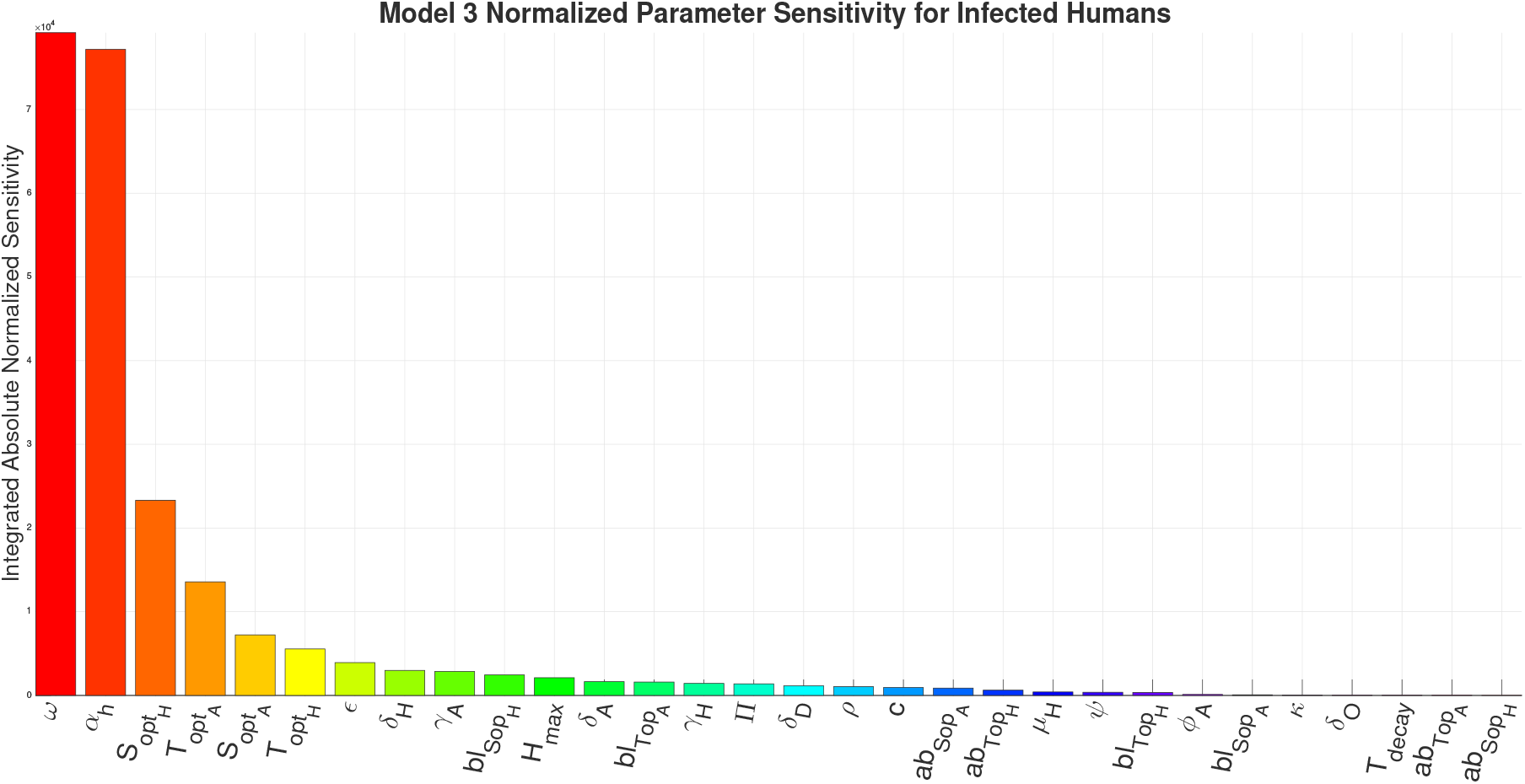
Model 3 Integrated Absolute Normalized Sensitivity analysis.

**Figure 16:**
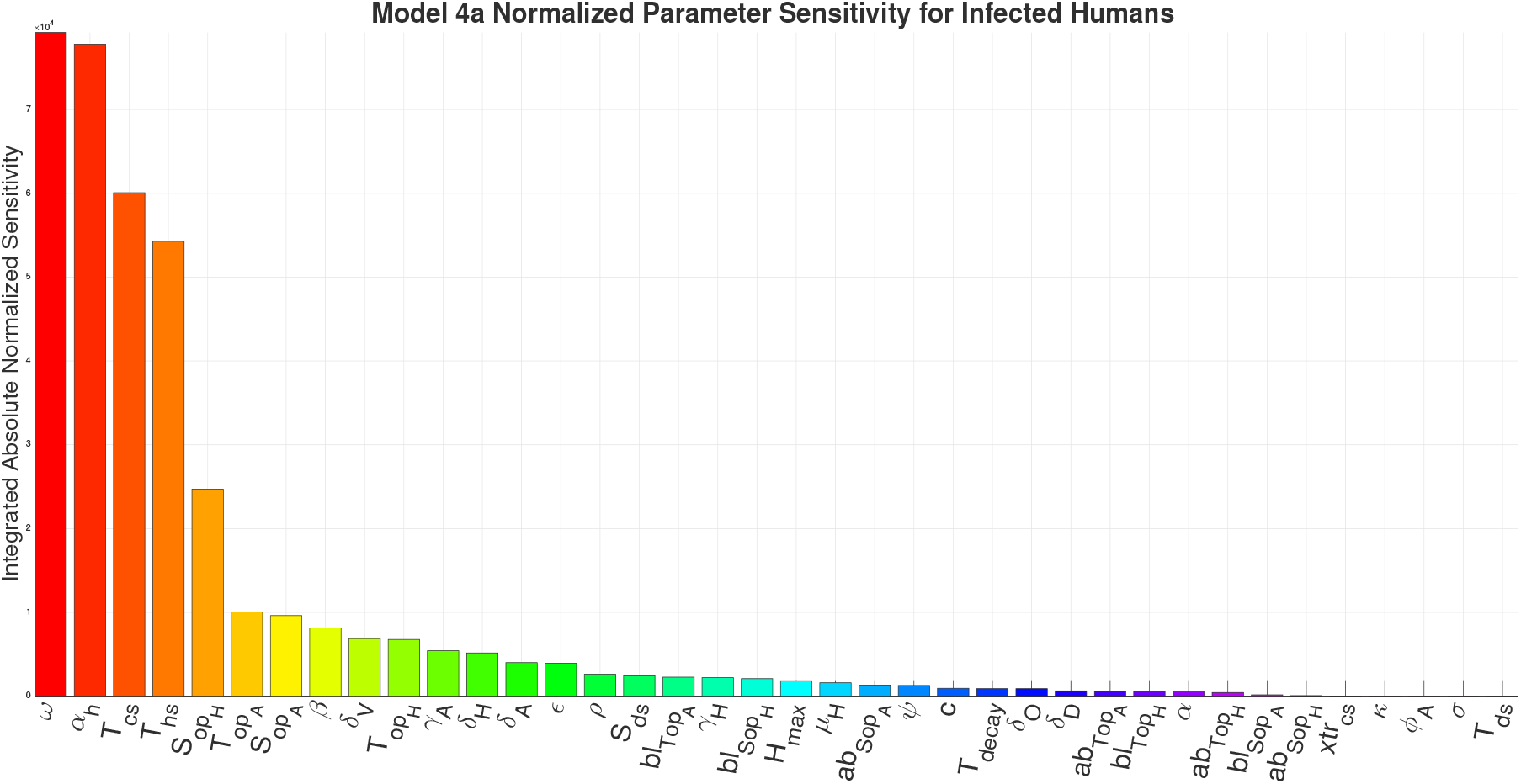
Model 4a Integrated Absolute Normalized Sensitivity analysis.

**Figure 17:**
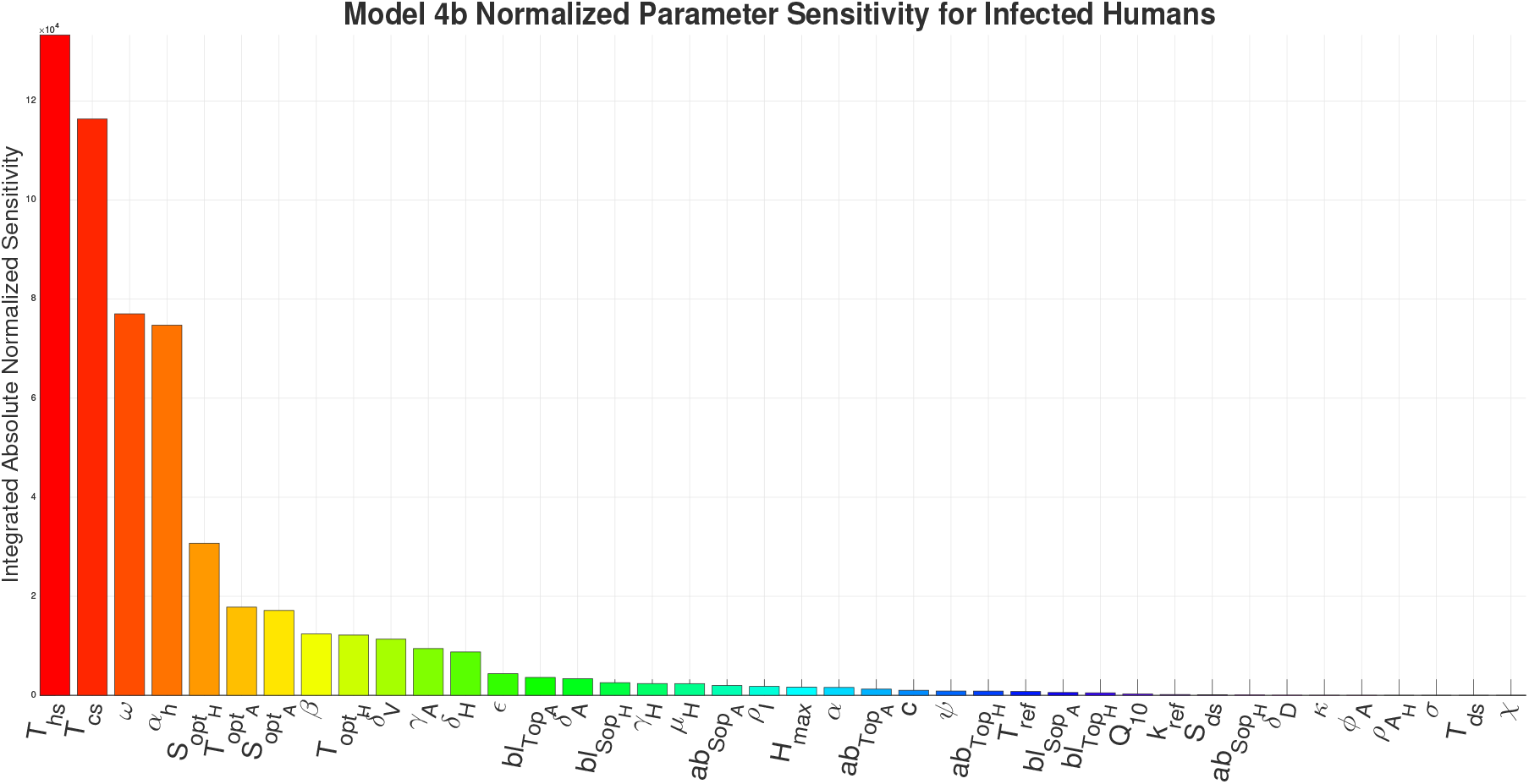
Model 4b Integrated Absolute Normalized Sensitivity analysis.

## 5 Reproduction Number

In the Valley fever sapronotic system for all the models we have laid out (Models 1–4b), biologically the basic reproduction number (*R*_0_) is represents the expected number of new hyphae or arthroconidia, or units of biomass, produced by a single introduced hyphae or arthroconidia unit in a naive environment. Because of this, *R*_0_ would be more precisely called the environmental reproduction number, which we will instead define as *R*_*E*_ to avoid confusion with *R*_0_ as defined for diseases spread from human to human. The term *R*_*E*_ determines whether the fungus survives, not whether humans get infected. If *R*_*E*_ > 1, *Coccidioides* becomes endemic and grows in the environment, creating sustained exposure risk for humans. If *R*_*E*_ < 1, *Coccidioides* dies out in the environment, and human cases drop to zero because the source is gone.

In all the models, the human disease compartments function as spillover targets, which are defined as populations that acquire infection from the environmental reservoir but act as epidemiological “dead ends” that do not increase the amount of pathogen or return it to the environment, thereby not contributing to the maintenance of the environmental reproduction number. In Model 1, the Infected compartment (*I*) is the spillover target. In Models 2, 3, and 4a, the Exposed (*E*) and Infected (*I*) compartments are the spillover targets. In Model 4b, in addition to *E* and *I*, the Asymptomatic compartment (*A*_*H*_) is a spillover target. The Wildlife compartment (*W*) in Models 4a and 4b serves as a reservoir host for the fungus, while humans remain dead-end hosts. Because humans act as dead-end hosts and do not contribute to fungal growth, human compartments do not appear in the infection subsystem that defines the next-generation matrix. Thus, *R*_*E*_ is determined entirely by the fungal subsystem. Although *R*_*E*_ governs fungal persistence rather than direct human-to-human transmission, sustained values of *R*_*E*_ ¿1 imply a persistent environmental reservoir, which, in turn, maintains long-term human exposure and endemic disease incidence.

All the models can be used to gain qualitative insight into the dynamics of Coccidioidomycosis, and for Models 3–4b, to understand how the environment influences the spread. But to get a deeper understanding of the underlying dynamics of all the models, we need to explore some basic qualitative properties. For each respective model, we identify the feasible region for the model, find the disease-free equilibrium (DFE), and then use the next-generation method [72, 73, 74, 75] to determine the basic reproduction number, *R*_*E*_. For models incorporating climate-dependent forcing (Models 3–4b), the resulting *R*_*E*_ (*T, S*_*m*_) is time-dependent through environmental drivers. Strictly speaking, this corresponds to an instantaneous or environment-dependent reproduction number rather than a classical constant threshold. However, it still provides a meaningful local criterion for fungal persistence under fixed environmental conditions. The full step-by-step breakdown to find *R*_*E*_ and defining the feasible region, proving the system is well-posed, with solutions remaining non-negative and bounded within a positively invariant region for each model can be found in the Supplementary Information. We will only outline the *F, V*, and resulting *R*_*E*_ in this section.

### 5.1 Model 1 Reproduction Number

The Model 1 disease-free equilibrium was found by setting the right-hand sides of the Equations 1-5 and the associated infected compartments (*H, I*) of Model 1, to zero, and solving for the state variables in the remaining compartments for the rate of change to be zero. This process yields, 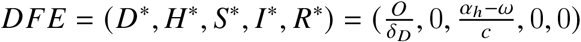. Humans do not infect susceptible humans or return fungus to the soil in this model.Then we construct the transmission matrix *F*,

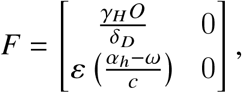

and we construct the transition matrix *V*,

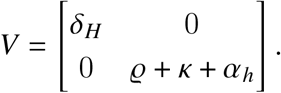

From matrix *F* and *V* we find the spectral radius of *F* · *V*^−1^ to yield *R*_*E*_,

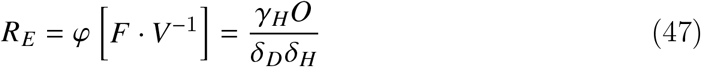

for Model 1.

### 5.2 Model 2 Reproduction Number

The infected compartments of Model 2 are set as (*H, A, E, I*), where *H* and *A* are infected fungal compartments and *E* and *I* are infected human compartments. The Model 2 disease-free equilibrium was found by setting the right-hand sides of the Equations 7-15 and the associated infected compartments (*H, A, E, I*) of Model 2, to zero, and solving for the state variables in the remaining compartments for the rate of change to be zero. This process yields, 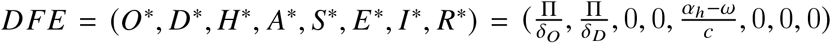. Then we construct the transmission matrix *F*,

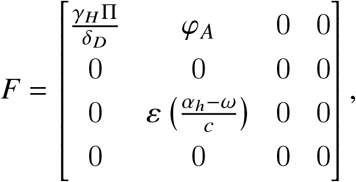

and we construct the transition matrix *V*,

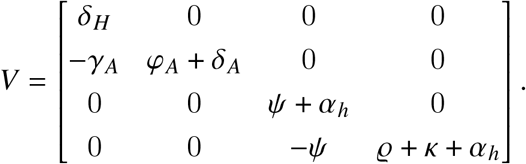

From matrix *F* and *V* we find the spectral radius of *F* · *V*^−1^ to yield *R*_*E*_,

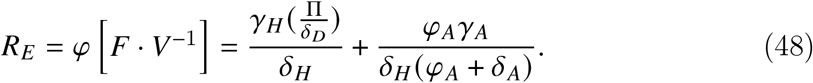

### 5.3 Model 3 Reproduction Number

The infected compartments of Model 3 are set as (*H, A, E, I*), where *H* and *A* are infected fungal compartments and *E* and *I* are infected human compartments. The Model 3 disease-free equilibrium was found by setting the right-hand sides of the Equations 20-23, 12-15, and the associated infected compartments (*H, A, E, I*) of Model 3, to zero, and solving for the state variables in the remaining compartments for the rate of change to be zero. This process yields the disease-free equilibrium, 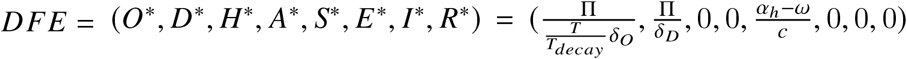. Then we construct the transmission matrix *F*,

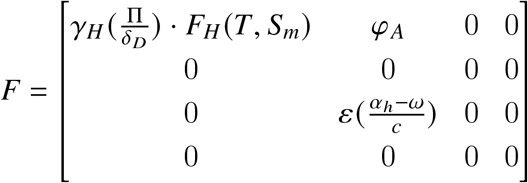

and we construct the transition matrix *V*,

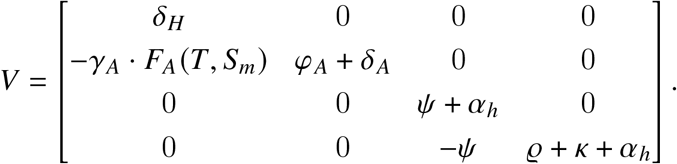

From matrix *F* and *V* we find the spectral radius of *F* · *V*^−1^ to yield *R*_*E*_,

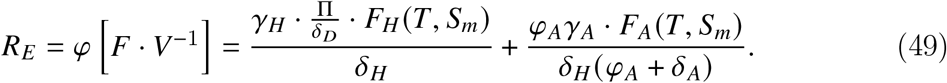

### 5.4 Model 4a Reproduction Number

The infected compartments of Model 4a are set as (*H, A, E, I*), where *H* and *A* are infected fungal compartments and *E* and *I* are infected human compartments. The Model 4a disease-free equilibrium was found by setting the right-hand sides of the Equations 25-29, 12-15, and the associated infected compartments (*H, A, E, I*) of Model 4a, to zero, and solving for the state variables in the remaining compartments for the rate of change to be zero. This process yields two distinct DFEs, a “wildlife-free” equilibrium 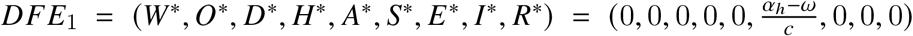 and a “wildlife-present” equilibrium 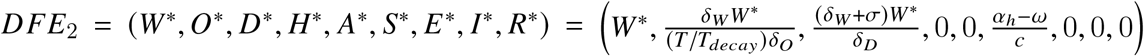. The “wildlife-free” DFE is not biologically plausible but the correlated *R*_*E*_ can be found in the Supplementary Information.

Then for the “wildlife-present” equilibrium *DFE*_2_ we construct the transmission matrix *F*,

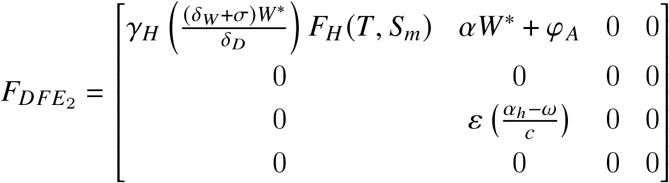

and the transition matrix *V*,

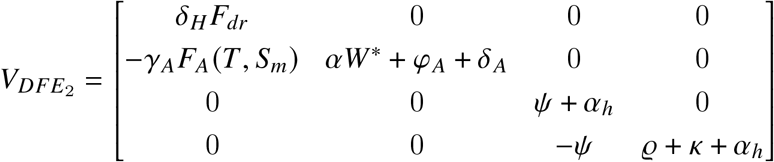

Then from matrix *F* and *V* we find the spectral radius of [*FV*^−1^] to yield *R*_*E*_,

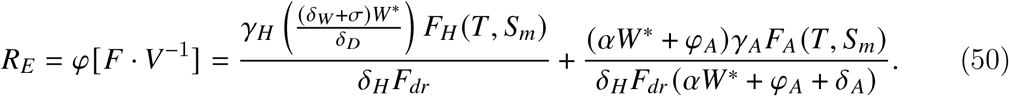

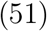

### 5.5 Model 4b Reproduction Number

The infected compartments of Model 4b are set as (*H, A, E, A*_*H*_, *I*), where *H* and *A* are infected fungal compartments and *E, A*_*H*_, and *I* are infected human compartments. The Model 4b disease-free equilibrium was found by setting the right-hand sides of the Equations 32-36, 37-41, and the associated infected compartments (*H, A, E, A*_*H*_, *I*) of Model 4b, to zero, and solving for the state variables in the remaining compartments for the rate of change to be zero. This process yields two distinct DFEs, a “wildlife-free” equilibrium 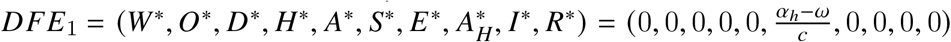 and a wildlife-present” equilibrium 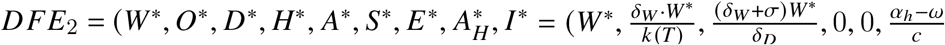. The “wildlife-free” DFE is not biologically plausible but the correlated *R*_*E*_ can be found in the Supplementary Information.

Then for the “wildlife-present” equilibrium *DFE*_2_ we construct the transmission matrix *F*,

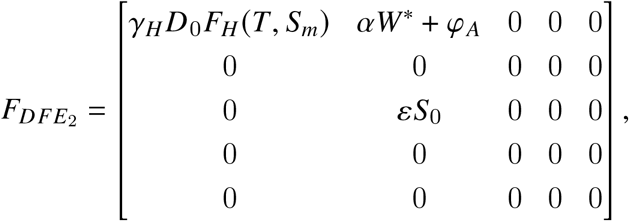

and the transition matrix *V*,

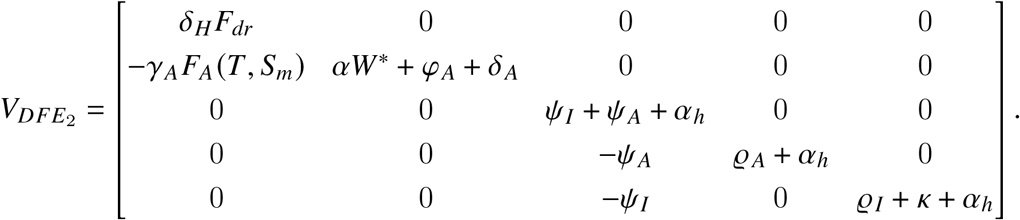

Then, from matrix *F* and *V* we find the spectral radius of [*FV*^−1^] to yield *R*_*E*_,

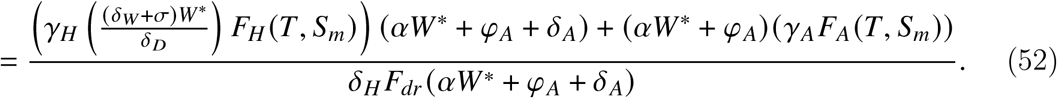

### 5.6 Effect of Climate and Wildlife on Reproduction Numbers

In Models 3, 4a, and 4b the reproduction number involved functions that depend on temperature and soil moisture. It should be noted that *R*_*E*_ strictly measures environmental hazard, which are the climatic and ecological conditions that allow the fungus to persist and multiply in the soil, rather than direct human risk. In models 4a and 4b there are also wildlife population terms (*W*^*^) in the respective derived *R*_*E*_ values. We set values for all parameters to the Maricopa County fit values for each model. Then we create heat maps showing how the value of *R*_*E*_ changes with respect to temperature and soil moisture. Models 4a and 4b have various subfigures showing how *R*_*E*_ changes with respect to temperature and soil moisture and different wildlife population levels. The maximum/minimum temperature recorded in Maricopa County is 16°F/122°F. The maximum/minimum soil moisture (palmer z-index) level recorded in Maricopa County is 3.28/10.77. The ranges were extended slightly to 15°F/125°F and -4/13 respectively.

Figure 18 visualizes the climatic niche for Coccidioides persistence as defined by Model 3. The *R*_*E*_ value is entirely dependent on the multiplicative environmental functions *F*_*H*_ (*T, S*_*m*_) and *F*_*A*_ (*T, S*_*m*_). The figure shows a single, contiguous “hot zone” where *R*_*E*_ > 1, peaking in the dark-red core where *R*_*E*_ attains its highest values under optimal environmental conditions. This peak occurs in a range of approximately 55-85°F and a soil moisture (Palmer Z-index) between 2 and 8. The shape is dictated by the black lines representing the fitted optimal climate parameters. The *R*_*E*_ value drops off sharply below the optimal soil moisture for hyphae 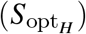, indicating that this parameter creates a critical threshold for fungal viability in this model. As this model lacks the influence of wildlife, this plot represents a baseline environmental potential for the fungus, driven only by climate.

**Figure 18:**
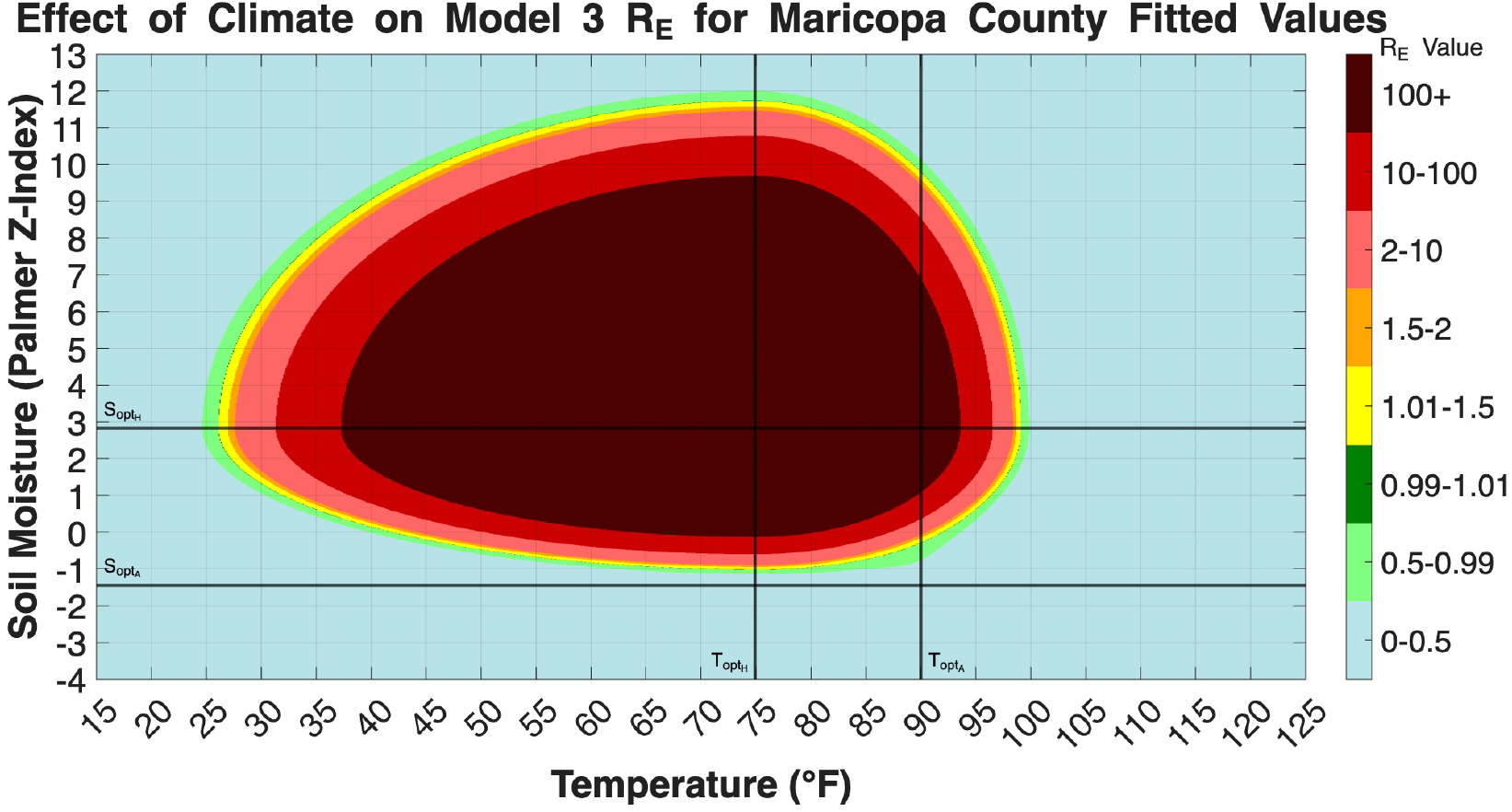
Heatmap of the *R*_*E*_ for Model 3, using parameters fit to Maricopa County data. The *R*_*E*_ value (color scale) is plotted as a function of Temperature (x-axis) and Soil Moisture (Palmer Z-Index, y-axis). The vertical black lines indicate the optimal fit temperatures for hyphae 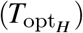 and arthroconidia 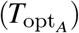 growth. The horizontal black lines indicate the optimal fit soil moisture for hyphae 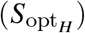 and arthroconidia 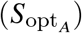 growth. The *R*_*E*_ value is calculated according to Equation 49, which is a function of the environmental drivers *F*_*H*_ (*T, S*_*m*_) and *F*_*A*_ (*T, S*_*m*_) but does not include wildlife dynamics. The region where *R*_*E*_ > 1 (yellow to dark red) represents the climatic conditions under which Coccidioides can potentially persist endemically.

Figure 19 shows how big of an impact the wildlife population, *W*^*^, has on the *R*_*E*_ for Model 4a. The *R*_*E*_ equation for this model includes *W*^*^ in the numerator of both its primary terms, representing wildlife’s dual role in providing organic substrate from death and spreading spores from activity. The four subplots clearly illustrate the dependency, as at a very low wildlife population (*W*^*^ = 2), the region where *R*_*E*_ > 1 is exceptionally small, with a peak *R*_*E*_ barely exceeding 1.5. As the wildlife population increases to *W*^*^ = 25 and *W*^*^ = 250, the region where *R*_*E*_ > 1 expands significantly, and the peak *R*_*E*_ values increase. Finally, at a high wildlife population of *W*^*^ = 10000, the environmental conditions for endemic persistence (*R*_*E*_ > 1) cover a massive region of the plot, with a large core where *R*_*E*_ > 100. This visualization is consistent with the sensitivity analysis, indicating that wildlife population levels act as a strong amplifying factor in the environmental transmission cycle.

**Figure 19:**
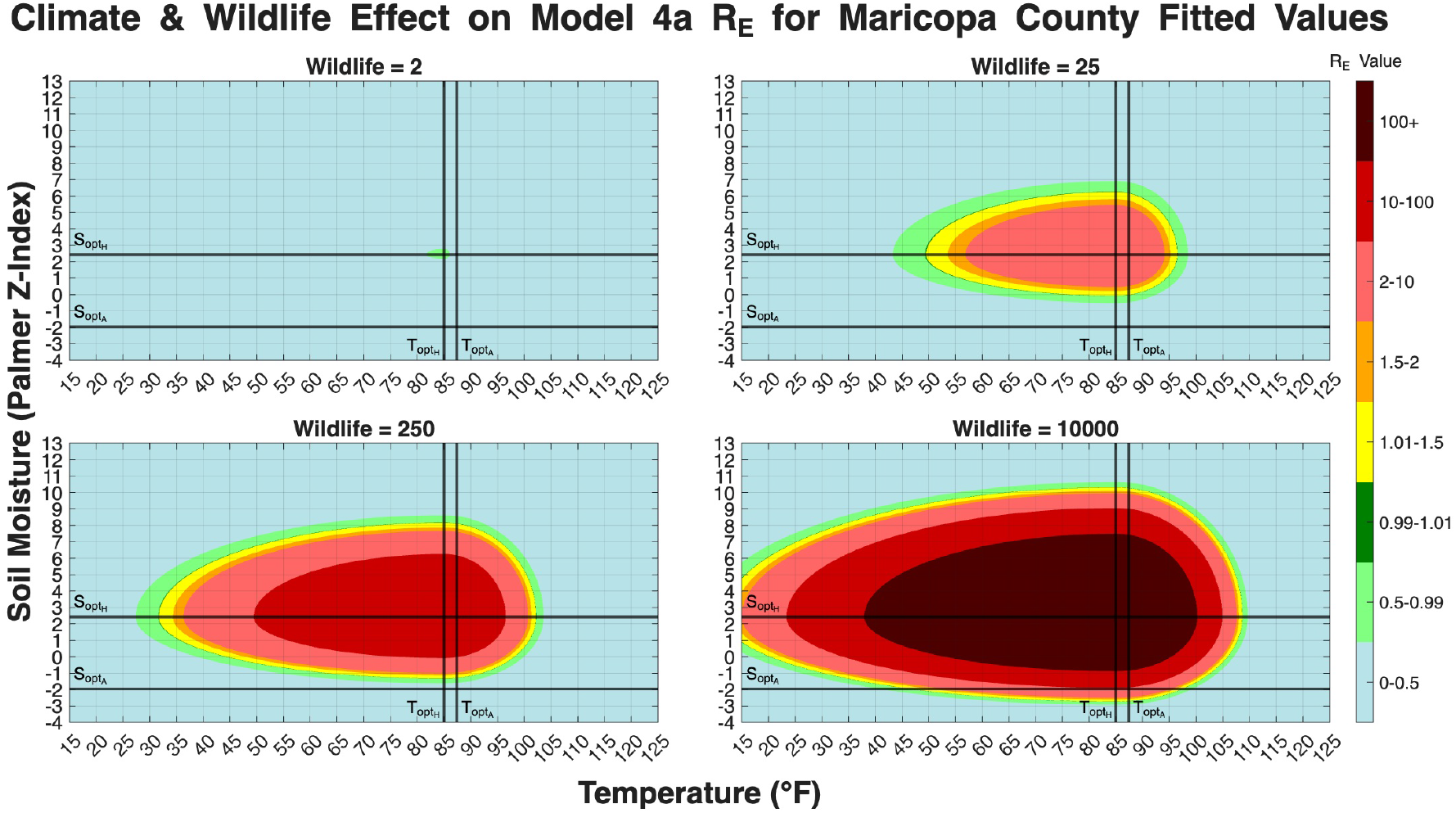
Heatmaps of *R*_*E*_ for Model 4a, using parameters fit to Maricopa County data. The figure has four subplots, each corresponding to a different fixed value for the wildlife population equilibrium (*W*^*^): 2, 25, 250, and 10000. The *R*_*E*_ value represented as a color scale is plotted against Temperature, in the x-axis, and Soil Moisture represented by Palmer Z-index, in the y-axis. The *R*_*E*_ calculation is defined by Equation 50, which incorporates *W*^*^ as a scaling factor for both the hyphae and arthroconidia growth terms. The black lines represent the fitted optimal climate parameters for hyphae and arthroconidia.

Figure 20, plots *R*_*E*_ for Model 4b, shows a dynamic that is visually almost identical to that of Model 4a in Figure 19. As with Model 4a, the wildlife population (*W*^*^) is the dominant factor controlling the size and intensity of the *R*_*E*_ > 1 region. This visual similarity suggests that while Model 4b provided a statistically better fit to the human case data, as seen in Table 1, the relationship between climate, wildlife, and *R*_*E*_ is not substantially altered by the more mechanistic *Q*_10_ decay function compared to the simpler temperature-dependent decay in Model 4a. This figure reinforces the conclusion that the inclusion of the wildlife compartment is the most significant structural enhancement for capturing the disease’s potential, with its population level acting as a primary amplifier for the entire environmental transmission cycle.

**Figure 20:**
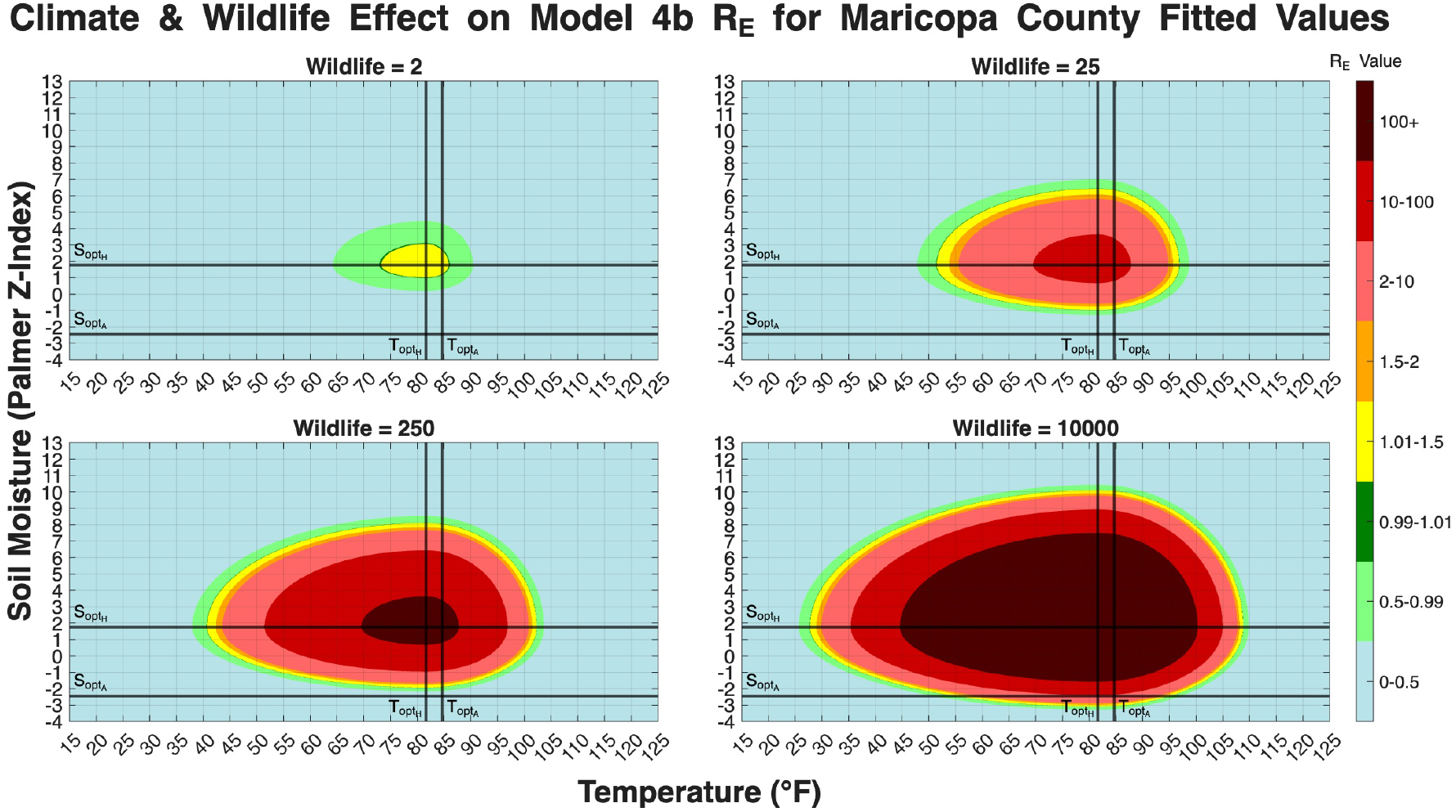
Heatmaps of *R*_*E*_ for Model 4b, using parameters fit to Maricopa County data. The *R*_*E*_ value represented as a color scale is plotted against Temperature, in the x-axis, and Soil Moisture represented by Palmer Z-index, in the y-axis. There are four fixed wildlife population levels (*W*^*^ = 2, 25, 250, and 10000). Model 4b differs from 4a by incorporating a more mechanistic *Q*_10_ model for organic decay and an asymptomatic human compartment. The *R*_*E*_ value is calculated according to Equation 52. The black lines represent the fitted optimal climate parameters.

## 6 Discussion and Conclusions

We developed a progressive series of five mechanistic models to unravel the transmission dynamics of Coccidioidomycosis (Valley fever). One of the main goals of this research was to build a flexible mechanistic framework that integrates environmental, fungal, and zoological drivers of the disease’s etiology, with the potential to be adapted to other endemic regions. By beginning with a basic fungal growth model and subsequently introducing additional layers of complexity, such as environmental drivers, wildlife, and more nuanced biological interactions, we have developed a structured modeling framework for investigating the determinants of the prevalence of *Coccidioides* in the environment and the production of human cases.

The results of our analyses indicate that including additional biologically motivated mechanisms, particularly environmental forcing and wildlife dynamics, substantially improves the models’ ability to reproduce observed temporal patterns in human case data. The initial models (Model 1 and Model 2) established a baseline but struggled to account for seasonal fluctuations. The incorporation of temperature and soil moisture dependencies in Model 3 represented a substantial advancement, highlighting the pivotal role of climate in the fungal saprobic cycle. However, it was the introduction of a wildlife compartment in Model 4a and a more sophisticated

*Q*_10_ representation of organic decay in Model 4b that resulted in the lowest RRMSE values, both in fitting and forecasting, in all assessed regions. Another important note is that consistently lower AIC, AICc, and BIC values relative to simpler models, such as the statistical Tamerius/Comrie Model or Model 1 and 2, indicate an improved balance between goodness-of-fit and model complexity. Model 4b achieved the most optimal balance of goodness-of-fit and model parsimony.

A challenge we want to emphasize is how, in Valley fever epidemiology, phenomenological statistical models often do not capture extreme outbreak peaks. Through the application of expanding-window forecasting and absolute loss differentials (DM and MDM tests), a clear, progressive trajectory of predictive power was quantified across the model hierarchy. Model 2, which incorporated fungal life-cycle stages without environmental constraints, performed statistically worse than the phenomenological baseline (MDM = -2.0491, *p* = 0.0423). The poor performance of Model 2 suggests that modeling fungal growth without incorporating climate-driven dynamics is insufficient to capture observed outbreak variability.

Incorporating temperature and soil moisture in Model 3 improved forecasting performance, eliminating the statistical baseline’s advantage (no significant difference) and establishing climate as a key data set in mechanistic modeling. Models 4a and 4b demonstrated statistically significant predictive superiority over the baseline (MDM=1.2353/*p* = 0.0264 and MDM=1.8613/*p* = 0.0012, respectively). The progression of these models gives us a clear indication that while climate determines the baseline transmission potential for *Coccidioides*, integrating mechanistic amplification through a wildlife reservoir substantially improves the ability to forecast severe real-world outbreaks.

The sensitivity analysis yielded valuable insights into the parameters that exert the most significant influence on disease outcomes. Across all models, parameters related to fundamental human population dynamics, *α*_*h*_ and *ω*, demonstrated consistent high sensitivity, as expected in an open-population framework. Interestingly, as the models became more complex, the parameters governing the fungal life cycle (*γ*_*H*_, *F*_*H*_, *F*_*A*_) and the reproductive phenology of wildlife (*T*_*hs*_, *T*_*cs*_) became more sensitive. The high sensitivity of wildlife heat- and cold-stress parameters indicates that wildlife demographic processes significantly influence fungal transmission dynamics within Models 4a and 4b. This finding underscores the importance of accurately characterizing the interconnected ecological niche of *Coccidioides* to produce reliable predictions.

The derivation of the environmental reproduction number (*R*_*E*_) in Section 5.6 further reinforces these conclusions from a mechanistic perspective. The *R*_*E*_ heatmaps provide a clear visualization of the pathogen’s viability threshold. As shown in Figure 18, Model 3 illustrates the specific temperature and soil moisture conditions where *R*_*E*_ > 1. Figures 19 and 20 demonstrate that these climate conditions, when scaled by the wildlife reservoir population (*W*^*^), lead to substantial shifts in the modeled environmental persistence potential. At low wildlife levels (*W*^*^ = 2), the endemic threshold is barely reached, while at high levels (*W*^*^ = 10000), the *R*_*E*_ value and the corresponding endemic region expand massively. It is important to reiterate that *R*_*E*_ maps the environmental hazard (fungal persistence) rather than immediate human risk. Also, these results should be interpreted qualitatively, as the magnitude of *R*_*E*_ depends on model scaling and parameterization rather than a directly measurable biological quantity. Regardless, this finding provides a direct mathematical explanation for why the wildlife parameters emerged as highly sensitive in Model 4a and the most sensitive in Model 4b. This dynamic confirms that the reservoir population acts as the primary engine for the entire environmental saprobic cycle. All together, these results highlight that capturing the interaction between climate variability and ecological amplification mechanisms is critical for understanding the underlying dynamics and forecasting environmentally driven fungal diseases.

### 6.1 Future Work

It is important to acknowledge the limitations inherent in this study. A constraint we encountered was the scarcity of specific field and laboratory data about numerous fungal and wildlife related parameters. Consequently, these parameters were estimated by fitting the models to human case data. These fittings introduced practical identifiability challenges. Multiple parameter combinations across the environmental and wildlife compartments may yield similarly accurate epidemiological trajectories that do not fully capture true, localized biological values. Moreover, our representation of wildlife as a single, homogeneous compartment represents a simplification. The diverse ecologies, behaviors, and population cycles of different animals exert a more intricate influence on arthroconidia dispersal than the current models capture.

Future research should address these limitations by targeting data collection. Laboratory investigations to ascertain the optimal temperature and soil moisture conditions for *Coccidioides* growth would be instrumental in directly parameterizing the environmental functions (like *T*_*opt*_, *S*_*opt*_, *T*_*ds*_, *S*_*ds*_, *α*_*T*_, *α*_*S*_, *β*_*T*_, *β*_*S*_). Other parameters governing the physical fungal presence (*γ*_*H*_, *H*_*max*_, *δ*_*H*_, *γ*_*A*_, *δ*_*A*_, *φ*_*A*_) could be fit directly to longitudinal soil sampling data. Furthermore, specific organic decay tests in relevant arid climates would isolate the *Q*_10_-style temperature coefficient variables (*T*_*decay*_, *Q*_18_, *k*_*re f*_, *T*_*re f*_). Implementing an organic matter decay function with a lower decay rate at extreme temperatures may be relevant in Arizona regions. With reliable datasets of local wildlife population counts, the reservoir parameters (*β, T*_*hs*_, *T*_*cs*_, *δ*_*W*_) could be strictly bounded. This would also allow for the development of multispecies wildlife models incorporating varying home ranges and breeding patterns. Enhanced parameterization of these models would facilitate more reliable forecasting, thereby enabling public health officials to anticipate high-risk periods and implement targeted interventions, such as timely dust-mitigation mandates and enhanced clinical awareness campaigns. Specifically, accurate, early-warning forecasts generated by Model 4b could inform targeted public health strategies, such as targeted prophylactic measures for high-risk populations, preemptive alerts to local pulmonologists to reduce misdiagnosis rates during the early stages of symptomatic infection, and optimized allocation of antifungal therapeutics. Although the lack of empirical parameter datasets is a persistent issue in mechanistic modeling, we outline these criteria in the hope that future empirical studies can directly benefit and integrate with this mathematical framework.

As we push towards making a fully mechanistic model, the mechanism by which arthroconidia are aerosolized to the point that humans and other animals are exposed (e.g., construction, agricultural activities, earthquakes) is complex but is likely a meaningful addition. As with other compartments, getting granular, reliable data for these can be challenging. Lastly, as discussed in the Section 3.1, the human case data can fluctuate due to testing protocols and clinician awareness. Adding a function or specific compartments to handle time-varying testing protocols and clinician awareness explicitly could also be beneficial for forecasting efforts.

### 6.2 Conclusion

In conclusion, this paper introduces a novel, scalable framework for mechanistic modeling of Valley fever. By integrating key ecological and epidemiological components in small steps, we have demonstrated the value of a bottom-up approach to understanding this complex, environmentally driven disease. The strong forecasting performance of the more detailed models validates the critical role of the wildlife reservoir as an amplifier, demonstrating that spikes in wildlife breeding and subsequent die-offs drive colonization spread and nutrient windfalls required to fuel massive arthroconidia blooms. These findings support the continued development of mechanistic, environmentally coupled models as tools for understanding, forecasting, and public health planning of environmentally driven infectious diseases.

## Data Availability

Data and code used to produce model fitting, forecasting, and analyses used in this paper is available in the publicly accessible repository GitHub at https://github.com/trevorreckell/Mechanistic-Model-Valley-Fever.

## References

[1] C. Nguyen, B. M. Barker, S. Hoover, D. E. Nix, N. M. Ampel, J. A. Frelinger, M. J. Orbach, and J. N. Galgiani, “Recent advances in our understanding of the environmental, epidemiological, immunological, and clinical dimensions of coccidioidomycosis,” Clinical microbiology reviews, vol. 26, no. 3, pp. 505–525, 2013.

[2] K. Benedict et al., “Estimated burden of coccidioidomycosis,” JAMA Network Open, 2024. PMCID: PMC12134948.

[3] E. Rayens and K. Norris, “Economic burden of fungal diseases in the united states,” Medical Mycology, vol. 63, no. 6, p. myaf049, 2024.

[4] J. N. Galgiani, N. M. Ampel, J. E. Blair, A. Catanzaro, R. H. Johnson, D. A. Stevens, and P. L. Williams, “Coccidioidomycosis,” Clinical Infectious Diseases, vol. 41, no. 9, pp. 1217–1223, 2005.

[5] T. N. Kirkland, D. A. Stevens, C.-Y. Hung, S. Beyhan, J. W. Taylor, L. F. Shubitz, S. H. Duttke, A. Heidari, R. H. Johnson, S. C. Deresinski, et al., “Coccidioides species: A review of basic research: 2022,” Journal of Fungi, vol. 8, no. 8, p. 859, 2022.

[6] P. Ocampo-Chavira, R. Eaton-Gonzalez, and M. Riquelme, “Of mice and fungi: Coccidioides spp. distribution models,” Journal of Fungi, vol. 6, no. 4, p. 320, 2020.

[7] C. W. Emmons, “Coccidioidomycosis in wild rodents: A method of determining the extent of endemic areas,” Public Health Reports (1896-1970), vol. 58, no. 1, pp. 1–5, 1943.

[8] I. H. McHardy, B. M. Barker, and G. R. Thompson, “Review of clinical and laboratory diagnostics for coccidioidomycosis,” Journal of Clinical Microbiology, vol. 61, no. 5, p. e0158122, 2023.

[9] J. D. Tamerius and A. C. Comrie, “Coccidioidomycosis incidence in arizona predicted by seasonal precipitation,” PLoS One, vol. 6, no. 6, p. e21009, 2011.

[10] C. Anderson, Agent-based modeling of coccidioidomycosis. PhD thesis, University of Pittsburgh, 2013.

[11] J. W. Taylor and B. M. Barker, “The endozoan, small-mammal reservoir hypothesis and the life cycle of coccidioides species,” Medical Mycology, vol. 57, no. Supplement 1, pp. S16–S20, 2019.

[12] M. del Rocío Reyes-Montes, M.A. Pérez-Huitrón, J.L. Ocaña-Monroy, M.G. Frías-De-Leòn, E. Martínez-Herrera, R. Arenas, and E. Duarte-Escalante, “The habitat of coccidioides spp. and the role of animals as reservoirs and disseminators in nature,” BMC infectious diseases, vol. 16, no. 1, p. 550, 2016.

[13] N. A. Chow, D. Kangiser, L. Gade, O. Z. McCotter, S. Hurst, A. Salamone, R. Wohrle, W. Clifford, S. Kim, Z. Salah, et al., “Factors influencing distribution of coccidioides immitis in soil, washington state, 2016,” Msphere, vol. 6, no. 6, pp. e00598–21, 2021.

[14] R. R. Dobos, K. Benedict, B. R. Jackson, and O. Z. McCotter, “Using soil survey data to model potential coccidioides soil habitat and inform valley fever epidemiology,” PloS one, vol. 16, no. 2, p. e0247263, 2021.

[15] H. Du, P. Perré, et al., “A 3-variable pde model for predicting fungal growth derived from microscopic mechanisms,” Journal of Theoretical Biology, vol. 470, pp. 90–100, 2019.

[16] G. P. Boswell, H. Jacobs, F. A. Davidson, G. M. Gadd, and K. Ritz, “A positive numerical scheme for a mixed-type partial differential equation model for fungal growth,” Applied Mathematics and Computation, vol. 138, no. 2-3, pp. 321–340, 2003.

[17] M. Sautour, P. Dantigny, C. Divies, and M. Bensoussan, “A temperature-type model for describing the relationship between fungal growth and water activity,” International Journal of Food Microbiology, vol. 67, no. 1-2, pp. 63–69, 2001.

[18] A. C. Comrie, “Climate factors influencing coccidioidomycosis seasonality and outbreaks,” Environmental health perspectives, vol. 113, no. 6, pp. 688–692, 2005.

[19] K. Okuneye and A. B. Gumel, “Analysis of a temperature-and rainfall-dependent model for malaria transmission dynamics,” Mathematical biosciences, vol. 287, pp. 72–92, 2017.

[20] S. E. Eikenberry and A. B. Gumel, “Mathematical modeling of climate change and malaria transmission dynamics: a historical review,” Journal of mathematical biology, vol. 77, no. 4, pp. 857–933, 2018.

[21] D. M. Engelthaler, C. C. Roe, C. M. Hepp, M. Teixeira, E. M. Driebe, J. M. Schupp, L. Gade, V. Waddell, K. Komatsu, E. Arathoon, et al., “Local population structure and patterns of western hemisphere dispersal for coccidioides spp., the fungal cause of valley fever,” MBio, vol. 7, no. 2, pp. 10–1128, 2016.

[22] A. Kinlaw, “A review of burrowing by semi-fossorial vertebrates in arid environments,” Journal of Arid Environments, vol. 41, no. 2, pp. 127–145, 1999.

[23] D. M. Engelthaler, J. C. Chatters, and A. Casadevall, “Was coccidioides a pre-columbian hitchhiker to southcentral washington?,” MBio, vol. 14, no. 2, pp. e00232–23, 2023.

[24] P. C. Escherich, Social biology of the bushy-tailed woodrat, Neotoma cinerea, vol. 110. Univ of California Press, 1981.

[25] P.-F. Verhulst, “Notice sur la loi que la population suit dans son accroissement,” Correspondance mathématique et physique, vol. 10, pp. 113–121, 1838.

[26] A. T. Classen, D. E. Taylor, L. Forlini, A. Saini, and G. S. Sayler, “A logistic model of subsurface growth with application to bioremediation,” Bioremediation Journal, vol. 4, no. 1, pp. 11–20, 2000.

[27] W. O. Kermack and A. G. McKendrick, “A contribution to the mathematical theory of epidemics,” Proceedings of the royal society of london. Series A, Containing papers of a mathematical and physical character, vol. 115, no. 772, pp. 700–721, 1927.

[28] W. O. Kermack and A. G. McKendrick, “Contributions to the mathematical theory of epidemics. ii.–the problem of endemicity,” Proceedings of the Royal Society of London. Series A, Containing Papers of a Mathematical and Physical Character, vol. 138, no. 834, pp. 55–83, 1932.

[29] W. O. Kermack and A. G. McKendrick, “Contributions to the mathematical theory of epidemics. iii.–further studies of the problem of endemicity,” Proceedings of the Royal Society of London. Series A, Containing Papers of a Mathematical and Physical Character, vol. 141, no. 843, pp. 94–122, 1933.

[30] L. Edelstein-Keshet, Mathematical models in biology, Chpater 6. SIAM, 2005.

[31] M. Kot, Elements of mathematical ecology, Chapter 2. Cambridge University Press, 2001.

[32] J. E. Blair, N. M. Ampel, and S. E. Hoover, “Coccidioidomycosis in selected immunosuppressed hosts,” Medical mycology, vol. 57, no. Supplement 1, pp. S56–S63, 2019.

[33] A. Catanzaro, “Coccidioidomycosis,” in Seminars in Respiratory and Critical Care Medicine, vol. 25, pp. 123–128, Copyright© 2004 by Thieme Medical Publishers, Inc., 333 Seventh Avenue, New …, 2004.

[34] M. A. Mandel, B. M. Barker, S. Kroken, S. D. Rounsley, and M. J. Orbach, “Genomic and population analyses of the mating type loci in coccidioides species reveal evidence for sexual reproduction and gene acquisition,” Eukaryotic cell, vol. 6, no. 7, pp. 1189–1199, 2007.

[35] D. M. Engelthaler and S. A. Balajee, “Forensics and epidemiology of fungal pathogens,” in Microbial forensics, pp. 297–726, Elsevier, 2011.

[36] N. T. J. Bailey, The Mathematical Theory of Epidemics. London: Charles Griffin & Company Ltd, 1957.

[37] N. F. Crum, “Coccidioidomycosis: a contemporary review,” Infectious diseases and therapy, vol. 11, no. 2, pp. 713–742, 2022.

[38] V. Rossi, S. Giosué, and P. Racca, “A comparison of non-linear models for describing the effect of temperature on the development of plant pathogens,” European Journal of Plant Pathology, vol. 119, no. 2, pp. 157–168, 2007.

[39] A. S. Moffat, G. De-Pol, and A. D. Tomos, “The effect of temperature on the growth of aspergillus niger on a solid medium,” Mycological Research, vol. 99, no. 8, pp. 957–961, 1995.

[40] A. M. Gibson, J. Baranyi, and J. I. Pitt, “Predicting fungal growth: the effect of water activity on aspergillus flavus and related species,” International Journal of Food Microbiology, vol. 23, no. 3-4, pp. 419–431, 1994.

[41] N. Magan and J. Lacey, “Effect of temperature and ph on water relations of field and storage fungi,” Transactions of the British Mycological Society, vol. 82, no. 1, pp. 71–81, 1984.

[42] J. Leplat, H. Friberg, M. Abid, and C. Steinberg, “Survival of fusarium graminearum macroconidia in soil: development of a predictive model of saprophytic activity,” Fungal Ecology, vol. 24, pp. 60–69, 2016.

[43] M. del Rocío Reyes-Montes, M.A. Pérez-Huitrón, J.L. Ocaña-Monroy, M.G. Frías-De-Leòn, E. Martínez-Herrera, R. Arenas, and E. Duarte-Escalante, “The habitat of coccidioides spp. and the role of animals as reservoirs and disseminators in nature,” BMC infectious diseases, vol. 16, no. 1, pp. 1–9, 2016.

[44] The Center for Food Security and Public Health (CFSPH), “Coccidioidomycosis (valley fever),” Sep 2021. Iowa State University, College of Veterinary Medicine. Accessed: November 4, 2025.

[45] Valley Fever Center for Excellence (VFCE), “Valley fever in other animals,” N.D. University of Arizona. Accessed: November 4, 2025.

[46] Merck Veterinary Manual, “Coccidioidomycosis in animals,” Oct 2022. Accessed: November 4, 2025.

[47] F. H. Bronson, “Climate change and seasonal reproduction in mammals,” Philosophical Transactions of the Royal Society B: Biological Sciences, vol. 364, no. 1534, pp. 3331–3340, 2009.

[48] R. Barrientos, A. Barbosa, F. Valera, and E. Moreno, “Temperature but not rainfall influences timing of breeding in a desert bird, the trumpeter finch (bucanetes githagineus),” Journal of Ornithology, vol. 148, pp. 411–415, 2007.

[49] R. M. Chew and A. E. Chew, “Energy relationships of the mammals of a desert shrub (larrea tridentata) community,” Ecological Monographs, vol. 40, no. 1, pp. 1–21, 1970.

[50] J.H. van‘tHoff, Ètudes de dynamique chimique. Frederik Muller & Co., 1884.

[51] Q. Wu, R. Ye, S. D. Bridgham, and Q. Jin, “Limitations of the q10 coefficient for quantifying temperature sensitivity of anaerobic organic matter decomposition: a modeling based assessment,” Journal of Geophysical Research: Biogeosciences, vol. 126, no. 8, p. e2021JG006264, 2021.

[52] L. Friedman, C. E. Smith, D. Pappagianis, and R. Berman, “Survival of coccidioides immitis under controlled conditions of temperature and humidity,” American Journal of Public Health and the Nations Health, vol. 46, no. 10, pp. 1317–1324, 1956.

[53] H. L. Mead, P. S. Hamm, I. N. Shaffer, M. d. M. Teixeira, C. S. Wendel, N. P. Wiederhold, G. R. Thompson III, R. Muñiz-Salazar, L.R. Castañón-Olivares, P. Keim, et al., “Differential thermotolerance adaptation between species of coccidioides,” Journal of Fungi, vol. 6, no. 4, p. 366, 2020.

[54] M. E. Gorris, K. K. Treseder, C. S. Zender, and J. T. Randerson, “Coccidioidomycosis dynamics in relation to climate in the southwestern united states,” GeoHealth, vol. 2, no. 1, pp. 6–24, 2018.

[55] National Oceanic and Atmospheric Administration, “Palmer Drought Severity Index Explanation,” 2022.

[56] United States Environmental Protection Agency, “Download daily data,” 2026. Accessed: 2026-02-07.

[57] C. Tofallis, “A new method for regression and errors-in-variables based on minimizing the sum of squared relative errors,” South African Statistical Journal, vol. 43, no. 1, pp. 79–104, 2009.

[58] K. Morimune and K. Oya, “The superiority of r-squared as a measure of the goodness-of-fit in a heteroscedastic regression model,” Communications in Statistics-Theory and Methods, vol. 39, no. 10, pp. 1753–1764, 2010.

[59] W. S. Cleveland, Visualizing data. Hobart Press, 1993.

[60] T. Hastie, R. Tibshirani, and J. Friedman, The elements of statistical learning: data mining, inference, and prediction. Springer Science & Business Media, 2009.

[61] K. P. Burnham and D. R. Anderson, Model selection and multimodel inference: a practical information-theoretic approach. Springer-Verlag, 2002.

[62] H. Akaike, “A new look at the statistical model identification,” IEEE Transactions on Automatic Control, vol. 19, no. 6, pp. 716–723, 1974.

[63] G. Schwarz, “Estimating the dimension of a model,” The Annals of Statistics, vol. 6, no. 2, pp. 461–464, 1978.

[64] R. J. Hyndman and G. Athanasopoulos, Forecasting: principles and practice. OTexts, 3rd ed., 2021. https://otexts.com/fpp3/.

[65] L. J. Tashman, “Out-of-sample tests of forecasting accuracy: an analysis and review,” International Journal of Forecasting, vol. 16, no. 4, pp. 437–450, 2000.

[66] F. X. Diebold and R. S. Mariano, “Comparing predictive accuracy,” Journal of Business & Economic Statistics, vol. 13, no. 3, pp. 253–263, 1995.

[67] D. Harvey, S. Leybourne, and P. Newbold, “Testing the equality of prediction mean squared errors,” International Journal of Forecasting, vol. 13, no. 2, pp. 281–291, 1997.

[68] A. Saltelli, M. Ratto, T. Andres, F. Campolongo, J. Cariboni, D. Gatelli, M. Saisana, and S. Tarantola, Global Sensitivity Analysis: The Primer. John Wiley & Sons, 2008.

[69] A. Varma, M. Morbidelli, and H. Wu, Parametric Sensitivity in Chemical Systems. Cambridge University Press, 1999.

[70] T. Turányi, “Sensitivity analysis of complex kinetic systems,” Journal of Mathematical Chemistry, vol. 5, no. 3, pp. 203–248, 1990.

[71] B. P. Ingalls, Mathematical Modeling in Systems Biology: An Introduction. MIT Press, 2013.

[72] O. Diekmann, J. A. P. Heesterbeek, and J. A. J. Metz, “On the definition and the computation of the basic reproduction ratio r 0 in models for infectious diseases in heterogeneous populations,” Journal of mathematical biology, vol. 28, pp. 365–382, 1990.

[73] O. Diekmann and J. A. P. Heesterbeek, Mathematical epidemiology of infectious diseases: model building, analysis and interpretation, vol. 5. John Wiley & Sons, 2000.

[74] O. Diekmann, J. Heesterbeek, and M. G. Roberts, “The construction of next-generation matrices for compartmental epidemic models,” Journal of the royal society interface, vol. 7, no. 47, pp. 873–885, 2010.

[75] P. Van den Driessche and J. Watmough, “Reproduction numbers and subthreshold endemic equilibria for compartmental models of disease transmission,” Mathematical biosciences, vol. 180, no. 1-2, pp. 29–48, 2002.

